# Regulation of effector gene expression as concerted waves in *Leptosphaeria maculans*: a two-players game

**DOI:** 10.1101/2021.12.15.472773

**Authors:** C. Clairet, E.J. Gay, A. Porquier, F. Blaise, C.L. Marais, M.-H. Balesdent, T. Rouxel, J.L. Soyer, I. Fudal

## Abstract

During infection, plant pathogenic fungi secrete a set of molecules collectively known as effectors, involved in overcoming the host immune system and in disease establishment. Effector genes are concertedly expressed as waves all along plant pathogenic fungi lifecycle. However, little is known about how coordinated expression of effector genes is regulated. Since many effector genes are located in repeat-rich regions, the role of chromatin remodeling in the regulation of effector expression was recently investigated. In *Leptosphaeria maculans*, causing stem canker of oilseed rape, we established that the repressive histone modification H3K9me3 (trimethylation of Lysine 9 of Histone H3), deposited by the histone methyltransferase KMT1, was involved in the regulation of expression of genes highly expressed during infection, including effectors. Nevertheless, inactivation of *KMT1* did not induce expression of these genes at the same level as observed during infection of oilseed rape, suggesting that a second regulator, such as a transcription factor (TF), might be involved. Pf2, a TF belonging to the Zn2Cys6 fungal specific TF family, was described in several Dothideomycete species as essential for pathogenicity and effector gene expression. We identified the orthologue of Pf2 in *L. maculans*, LmPf2, and investigated the role of LmPf2 together with KMT1, by inactivating and over-expressing *LmPf2* in a wild type (WT) strain and a *Δkmt1* mutant. Functional analyses of the corresponding transformants highlighted an essential role of LmPf2 in the establishment of pathogenesis. Transcriptomic analyses during axenic growth showed that LmPf2 is involved in the control of effector gene expression. We observed an enhanced effect of the over-expression of *LmPf2* on effector gene expression in a *Δkmt1* background, suggesting an antagonist role between KMT1 and LmPf2.

## INTRODUCTION

During infection, plant pathogenic fungi secrete a set of molecules, collectively known as effectors, involved in overcoming the host immune defense system, nutrient uptake and, eventually, symptom development (Lo Presti *et al*., 2015). Effectors correspond to secondary metabolites, siRNA, and small secreted proteins (SSP); the latter being often cysteine-rich, with rare homology to other known proteins in the databases (Weiberg *et al*., 2013; Lo Presti *et al*., 2015; Collemare *et al*., 2019). Effector genes of filamentous plant pathogens are often located in transposable elements (TEs)-rich regions of the genomes, such as dispensable chromosomes or telomeres (Sánchez-Vallet *et al*., 2018). Transcriptomic data generated during different stages of plant infection highlighted concerted waves of effector gene expression over the course of infection, according to the infection structure, to the host-plant infected or to the species or strain studied (e.g. Sanchez-Vallet *et al*., 2018; Haueisen *et al*., 2019; Gay *et al*., 2021). Little is known about how coordinated expression of effector genes is regulated. Based on observation that many effector genes are located in TE-rich regions, and up-regulated during host penetration/infection, the role of chromatin remodeling in the regulation of effector expression was recently investigated. Analyses in plant-interacting fungi have shed light on two histone methyltransferases, KMT1 and KMT6, as involved in the regulation of effector gene expression (Chujo & Scott 2014; Soyer *et al*., 2014; Soyer *et al*., 2019; Meile *et al*. 2020; Zhang *et al*., 2021).

*Leptosphaeria maculans*, the causal agent of stem canker of oilseed rape (*Brassica napus*), displays a complex life cycle with alternating stages of saprophytism, asymptomatic growth and necrotrophy (Rouxel and Balesdent, 2005). During infection of oilseed rape, several waves of genes are expressed, including predicted effector genes (Gay *et al*., 2021). Of particular interest, a specific set of effector genes is expressed during the asymptomatic stages occurring on leaves, petioles and stems, including the 11 avirulence genes (*AvrLm*) identified so far, these genes all being located in TE-rich regions (*AvrLm1, AvrLm2, AvrLm3, AvrLm4-7, AvrLm5-9, AvrLm6, AvrLm10A, AvrLm10B, AvrLm11, AvrLm14, AvrLmS-Lep2*; Gout *et al*., 2006; Fudal *et al*., 2007; Parlange *et al*., 2009; Balesdent *et al*., 2013; van de Wouw *et al*., 2014; Plissonneau *et al*., 2016; Ghanbarnia *et al*., 2015, 2018; Petit-Houdenot *et al*., 2019; Neik *et al*., 2020; Degrave *et al*., 2021). The genome of *L. maculans* displays a bipartite structure with gene-rich regions and TE-rich regions, the latter representing one third of the genome (Rouxel *et al*., 2011). In a recent study, we generated a map of the distribution of three histone modifications, either associated with euchromatin and gene expression (H3K4me2, dimethylation of Lysine 4 of histone H3) or heterochromatin resulting in gene silencing (H3K9me3 and H3K27me3, trimethylation of Lysine 9 and Lysine 27 of Histone H3) during axenic growth (Soyer *et al*., 2021). We highlighted an enrichment of effector genes in domains associated with either H3K9me3 or H3K27me3. Integrative analysis of ChIP-seq data *in vitro* with transcriptomic analyses performed throughout the life cycle of *L. maculans* pinpointed that *L. maculans* genes over-expressed at any stage of *B. napus* infection are enriched in heterochromatin domains *in vitro* (Gay *et al*., 2021). In a previous study, silencing of *KMT1* had led to an over-expression of more than 30% of the genes located in TE-rich environments normally silenced in axenic culture in the WT strain, specifically effector genes (Soyer *et al*., 2014). ChIP-qPCR analyses showed that over-expression of at least two effector genes was associated with a decrease of the repressive histone modification H3K9me3 in the genomic environment of these genes (Soyer *et al*., 2014). Altogether, these analyses revealed that chromatin structure, via the dynamic of chromatin remodeling, was an important regulatory layer of effector genes up-regulated during host infection and located in TE-rich regions. Nevertheless, at least in *L. maculans*, inactivation of *KMT1*, while inducing expression of effector genes *in vitro*, did not induce expression of these genes at the same level as observed during infection of oilseed rape, suggesting that a second regulator, such as a transcription factor (TF), might be involved. Hence, Soyer *et al*. (2015) proposed a model of dual control of effector gene expression: chromatin condensation repressed effector gene expression *in vitro*; after chromatin loosening upon infection, one or several TFs could bind effector gene promoters resulting in their concerted expression. Synergic involvement of chromatin modifications and TF(s) in fungal effector gene regulation remains poorly understood and promising field of investigation.

In filamentous plant pathogens, only a few TFs influencing effector gene expression have been identified so far (see for review Tan and Oliver, 2017). Among them, AbPf2, a Zn2Cys6 TF, was first described in *Alternaria brassicicola*, in which it regulates, directly or indirectly, expression of 33 genes encoding secreted proteins including eight putative effectors (Cho *et al*., 2013). In *Parastagonospora nodorum*, PnPf2 positively regulates two necrotrophic effector genes, *SnToxA* and *SnTox3*, and the orthologue of *SnToxA, ToxA*, is regulated by PtrPf2 in *Pyrenophora tritici-repentis* (Rybak *et al*., 2017). In this species, a recent transcriptomic analysis comparing the WT and the *PnPf2* mutant during axenic culture and infection of wheat revealed an involvement of *PnPf2* in the regulation of twelve effector-encoding genes and of genes associated with plant cell wall degradation and nutrient assimilation (Jones *et al*., 2019).

Here, we identified the homologue of Pf2 in *L. maculans* and investigated its involvement in the regulation of effector gene expression following removal of H3K9me3. We hypothesized that removal of H3K9me3 in the genomic environment of effector genes was a pre-requisite for induction of their expression through the action of one, or several, TFs during infection. We inactivated *LmPf2* via CRISPR-Cas9 and over-expressed *LmPf2* in two different genetic backgrounds: a wild type strain and a strain in which *KMT1* was inactivated. We characterized the corresponding transformants for their growth, sporulation and pathogenicity. We performed a RNA-seq analysis in order to decipher the involvement of LmPf2 and KMT1 in gene regulation. We found out that KMT1 and LmPf2 are acting antagonistically to regulate expression of genes expressed during infection of oilseed rape, specifically A*vrLm* genes as well as other genes (including putative effectors) concertedly expressed during infection.

## MATERIALS AND METHODS

### Fungal culture

The reference isolates JN3 (v23.1.3; Rouxel *et al*., 2011) and JN2 (v23.1.2; Balesdent *et al*., 2011), corresponding to two sister progenies of opposite mating types, were used as hosts for genetic transformations (Balesdent *et al*., 2001). We also used a JN2 strain constitutively expressing *eGFP* (Sâsêk *et al*., 2012) which was crossed with our mutants in order to follow the colonization of oilseed rape by the mutants. Fungal cultures and conidia production were performed as previously described (Ansan-Melayah *et al*., 1995). For DNA/RNA extractions, mycelium was grown on V8-juice agar medium at 25°C in the dark for seven days and then plugs were transferred into 150 ml of static Fries liquid medium in 500 ml Roux flasks. Tissues were harvested after growing for seven days at 25°C.

### Pathogenicity and growth assays

Pathogenicity assays were performed on cotyledons of 10-day-old plantlets of *B. napus*, ES-Astrid. Cotyledons were inoculated with pycnidiospore suspensions as described previously (Gall *et al*., 1995). Plants were incubated in a growth chamber at 19/24°C (night/day) with a 16h photoperiod and 90% humidity. Symptoms were scored on 10-12 plants, with two biological replicates, 13 days post inoculation (dpi), using the IMASCORE rating scale comprising six infection classes (IC), where IC1 to IC3 correspond to various levels of resistance of the plant and IC4 to IC6 to susceptibility (Balesdent *et al*., 2001). Growth assays were performed by deposition of a 5 mm plug at the center of 90 mm Petri dishes (containing 20 ml of V8-juice agar medium or MMII medium). Radial growth was measured at nine days after incubation in a growth chamber (25°C) on four biological replicates and statistical analyses were performed using Kruskal-Wallis test (Guo *et al*., 2013).

### Vector construction and fungal transformation

Vectors pLAU2 and pLAU53 conferring respectively hygromycin and geneticin resistance were used to perform CRISPR-Cas9 gene inactivation, as described by Idnurm *et al*. (2017). DNA fragments coding for guide RNA (gRNA) which target genes of interest were designed using the CRISPOR prediction tool and *L. maculans* JN3 strain as reference genome (http://crispor.tefor.net/; **Table S1**; Dutreux *et al*., 2018). The gRNA were chosen not to match on any other genes. The DNA fragment coding for gRNA was amplified using primers MAI0309 and MAI0310 and then inserted into the *Xho*I site of plasmid pLAU53 using Gibson assembly (Silayeva and Barnes, 2018). Hence, plasmids Plau53-KMT1 and Plau53-LmPf2 were generated to inactivate *KMT1* and *LmPf2*.

Over-expression plasmids were obtained using the pBht2 vector conferring resistance to hygromycin (Mullins *et al*., 2001). *EF1α* promoter was amplified using EF1aProSac1F and EF1aProKpnIR primers and genomic DNA of the WT isolate as template, and then inserted into *Kpn*I-*Sac*I digested pBht2 to obtain the pBht2-prom*EF1α* vector. The *LmPf2* gene and its terminator were then amplified using overex_*LmPf2*_Gibs_F and overex_ *LmPf2*_Gibs_R primers and inserted in 3’ of the *EF1α* promoter in the *Hin*dIII digested pBht2-promEF1α vector using Gibson assembly to obtain the plasmid pBht2-overex*LmPf2* (**Table S1**).

The constructs were introduced into the *Agrobacterium tumefaciens* strain C58-pGV2260 by electroporation (1.5 kV, 200 Ω and 25 mF). *Agrobacterium tumefaciens* mediated transformation (ATMT) of *L. maculans* was performed as previously described (Gout *et al*., 2006). Transformants were plated on minimal medium complemented with geneticin (50 mg/l) for pLAU53-gRNA or hygromycin (50 mg/l) for pBht2-overex*LmPf2* and pLAU2-Cas9 and cefotaxime (250 mg/l). For the CRISPR-Cas9 gene inactivation, construct containing *Cas9* (pLAU2-Cas9) was first introduced into the WT strain, transformants were selected for hygromycin resistance and this strain was subsequently transformed with pLAU53-gRNA directed against *KMT1* or *LmPf2*. Transformants were selected for geneticin, hygromycin and cefotaxime resistance. Mutations in the targeted genes were checked in the transformants by PCR amplifying with specific primers (**Table S1**) and sequencing. For the *LmPf2* over-expression, pBht2-overex*LmPf2* was introduced into the WT strain and the *Δkmt1* mutant. Insertion of the overex*LmPf2* construction was checked in transformants by PCR amplification (**Table S1**).

### Fungal crosses

Purifications in order to eliminate *Cas9* gene and gRNA-encoding gene from the CRISPR-Cas9 mutants were performed by crossing mutants with a WT strain of opposite mating type (expressing or not *GFP*; **Table S2**). Crosses were performed as described by Balesdent *et al*. (2002). Progeny was harvested and plated on V8-juice agar medium. Mycelium was collected and DNA extracted. PCR and sequencing were performed with primers check_CRISPR_*LmPf2* or _*KMT1* to select progeny with the targeted CRISPR-Cas9 mutation but without *Cas9* and gRNA-encoding gene.

### DNA and RNA manipulation

Genomic DNA was extracted from conidia or from mycelium grown in Fries liquid culture with the DNAeasy 96 plant Kit (Qiagen S.A., Courtaboeuf, France). PCR amplifications were performed as previously described (Fudal *et al*., 2007). To identify mutation arising in the sequence of the gene targeted by the CRISPR-Cas9 strategy, sequencing was performed by Eurofins Genomics (Anzinger, Ebersberg, Germany; **Table S1**). Total RNA was extracted from mycelium grown for one week in Fries liquid medium, and from cotyledons of oilseed rape infected by *L. maculans* seven dpi as previously described (Fudal *et al*., 2007).

### Quantitative RT-PCR

Quantitative RT-PCR (qRT-PCR) were performed using a model CFX96 Real Time System (BIORAD; Hercules, CA, USA) and Absolute SYBR Green ROX dUTP Mix (ABgene, Courtaboeuf, France) as previously described (Fudal *et al*., 2007). For each condition tested, two different RNA extractions from two different biological samples and two reverse transcriptions for each biological replicate were performed. Primers used for qRT-PCR are described in **Table S1**. Ct values were analyzed as described by Muller *et al*. (2002) or using the 2^-ΔΔCt^ method (Livak and Schmittgen, 2001). *β-tubulin* was used as a constitutively expressed reference gene.

### Confocal microscopy and binocular observation

Cotyledons of oilseed rape infected by strains expressing *GFP* were observed at two different time points (5 and 7 dpi) using a DM5500B Leica TCS SPE laser scanning confocal microscope and a 20x HCX Fluotar Leica objective lens. GFP was excited at 488 nm and emission was captured with a 505– 530 nm broad-pass filter. The detector gain was set-up between 700 and 900. Calibrations of gain settings were performed with multiple control leaves and with a range of background fluorescence. All images represent at least four scans. Infected cotyledons were also observed at 8 and 13 dpi using a Leica MZ16F fluorescent binocular coupled with a Leica DFC300FX camera.

### Western Blot

Total proteins were extracted from 10-100 mg lyophilized mycelium. Mycelium was ground using beads and Mixer Mill MM 400 (Retsch, Éragny, France). Total proteins were extracted, mixed with leammli buffer 4x (Biorad, Hercules, USA) and migrated on a polyacrylamide gel as described by Petit-Houdenot *et al*. (2019). Proteins were transferred on a PVDF membrane according to the manufacturer’s protocol (Trans-Blot^®^ Turbo™ Rapid Transfer System, BIORAD, Les-Ulis, France) using a small protein transfer program. The PVDF membrane was incubated in TBS 1X containing 5% powder milk, 0.05% tween 20 for one hour to saturate membrane. The membrane was then incubated at 4°C over-night in TBS 1% containing 0.05% Tween 20, milk 1% and an anti-H3K9me3 antibody (1:5000; 39062 ActiveMotif, La Hulpe, Belgium Germany). PVDF membrane was washed as described by Petit-Houdenot *et al*. (2019) and was then incubated in TBS 1X + 0.05% Tween 20 + milk 1% + anti-Rabbit IgG II^R^ (goat anti-rabbit, Santa Cruz Biotechnology, Heidelberg, Allemagne) and washed as previously. Finally, membrane was incubated 1 min in 1 ml enzyme solution (Clarity™ Western ECL, BIORAD, Les-Ulis, France) and 1 ml Luminol/enhancer solution (Clarity™ Western ECL, BIORAD, Les-Ulis, France) and observed using ChemiDoc (ChemiDoc™ Imaging Systems, BIORAD, Les-Ulis, France).

### Gene annotation and domain prediction

The *Pf2* orthologue of *L. maculans* had been previously identified by Rybak *et al*. (2017). As it had been identified on a previous version of the *L. maculans* genome, we performed a new search, using the protein sequence of Pf2 from *P. nodorum* and the NCBI BLASTP program (Altschul *et al*., 1990). Functional domains were identified using Pfam (Finn *et al*., 2014; https://pfam.xfam.org/). Alignments were performed with COBALT (Papadopoulos and Agarwala, 2007).

### RNA-seq and statistical analysis

Eight different transformants (i.e., two mutants inactivated for *KMT1* or *LmPf2* and two transformants in which *LmPf2* was over-expressed in a WT background (WT_o*Pf2*) or a *Δkmt1* background (*Δkmt1*_o*Pf2*, **Table S3**) were grown in static Fries liquid medium during 7 days and harvested for RNA extraction. Two RNA extractions corresponding to two biological replicates were performed for each transformant. About 150 mg of mycelium was used per extraction. Libraries were prepared from all biological replicates, individually, according to the Illumina TruSeq protocol (Illumina, San Diego, CA, USA). Libraries including polyA enrichment were performed and sequenced using 150 pb paired-end strategy on an HiSeq2000 Illumina sequencer at the Genewiz sequencing facility (Leipzig, Germany) with an input of 2 µg total RNA. Quality of the reads was checked and improved using Trimmomatic (Bolger *et al*., 2014). The resulting reads were treated to remove adaptors and reads below 30 bp and filtered reads were mapped against the *L. maculans* genome (Dutreux *et al*., 2018) using STAR (Dobin *et al*., 2013) with default parameters. Read alignments were stored in SAM format, and indexing, sorting, and conversion to BAM format were performed using SAMtools v0.1.19 (Li *et al*., 2009). Genes with a number of reads > 15 in at least one condition were kept for statistical analysis. Differential expression analyses were made using R, version 3.0.2 (www.r-project.org) and the package EdgeR (Robinson *et al*., 2010). Genes with a log2 Fold Change ≤ -1.5 or ≥ 1.5 and an associated False Discovery Rate ≤ 0.05 were considered as differentially expressed (McCarthy *et al*., 2012).

### Gene Ontology enrichment analysis

Gene Ontology (GO) annotations of *L. maculans* genes were retrieved from Dutreux *et al*. (2018). Gene ontology term enrichment analysis of the differentially expressed genes (DEG) in our transformants was performed with the plug-in Biological networks Gene ontology (BinGo; v3.0.3) of the cytoscape software (Shannon *et al*., 2003). List of genes submitted to BINGO were considered as significantly enriched for a given GO term with an associated False Discovery Rate ≤ 0.01 for the biological processes. All statistical analyses were done in R, version 3.0.2 (www.r-project.org).

### RNA-seq and ChIP-seq datasets

To investigate expression of avirulence genes, *KMT1* and *LmPf2*, and *in planta* expression of the DEG in the transformants generated in this study, we used previously generated *in vitro* and *in planta* RNA-seq data (Gay *et al*., 2021). Infection of oilseed rape had been performed using a WT strain. We used RNA-seq data from i) cotyledons of cultivar Darmor-*bzh* sampled 2, 5, 7, 9, 12 and 15 dpi (corresponding to EBI accession numbers SAMEA6086549, SAMEA104153286 and SAMEA104153287); ii) petioles of cultivar Darmor-*bzh* sampled 7 and 14 dpi (EBI accession numbers SAMEA104153278, ERS4810043, ERS4810044, ERS4810045); iii) stems of cultivar Bristol sampled 14 and 28 dpi (EBI accession numbers SAMEA104153293, ERS4810064, ERS4810065, ERS4810066, ERS4810067, ERS4810068) and iv) axenic growth of a WT strain on V8-medium (EBI accession numbers ERS4810062 and ERS4810063). To analyze location of the DEG in the transformants generated in this study in domains associated either with euchromatin or heterochromatin, we used previously generated genome-wide chromatin map (Soyer *et al*., 2021), available under the GEO accession number GSE150127. We assessed the significant enrichment of the DEG in H3K4me2-, H3K9me3- or H3K27me3-domains as described in Soyer *et al*. (2021). Enrichment was considered significant with a *P value* < 0.05; Chi^2^ tests were done using R, version 3.0.2 (www.r-project.org).

## RESULTS

### Identification of a *Pf2* orthologue and analysis of *KMT1* and *LmPf2* expression in *Leptosphaeria maculans*

In order to analyze involvement of Pf2 in the regulation of *L. maculans* gene expression, we first identified the *Pf2* orthologue in *L. maculans*. Phylogenetic analysis showed that the closest orthologue of *Pf2* of *P. nodorum* was found in *L. maculans* (Genbank Accession XP_003838593.1). In the recent *L. maculans* genome reannotation, this Genbank Accession corresponded to gene ID Lmb_jn3_06039, located on SuperContig 6 (Dutreux *et al*., 2018). A bidirectional Best Hit with BLASTp confirmed orthology between *PnPf2* and Lmb_jn3_06039 (**Figure 1**). Lmb_jn3_06039 (hereinafter referred to as *LmPf2*) encodes a 658 amino acids protein sharing 67% identity with PnPf2 with a Zn2Cys6-type DNA-binding domain (IPR001138; **Figure 1**).

**Figure 1:**
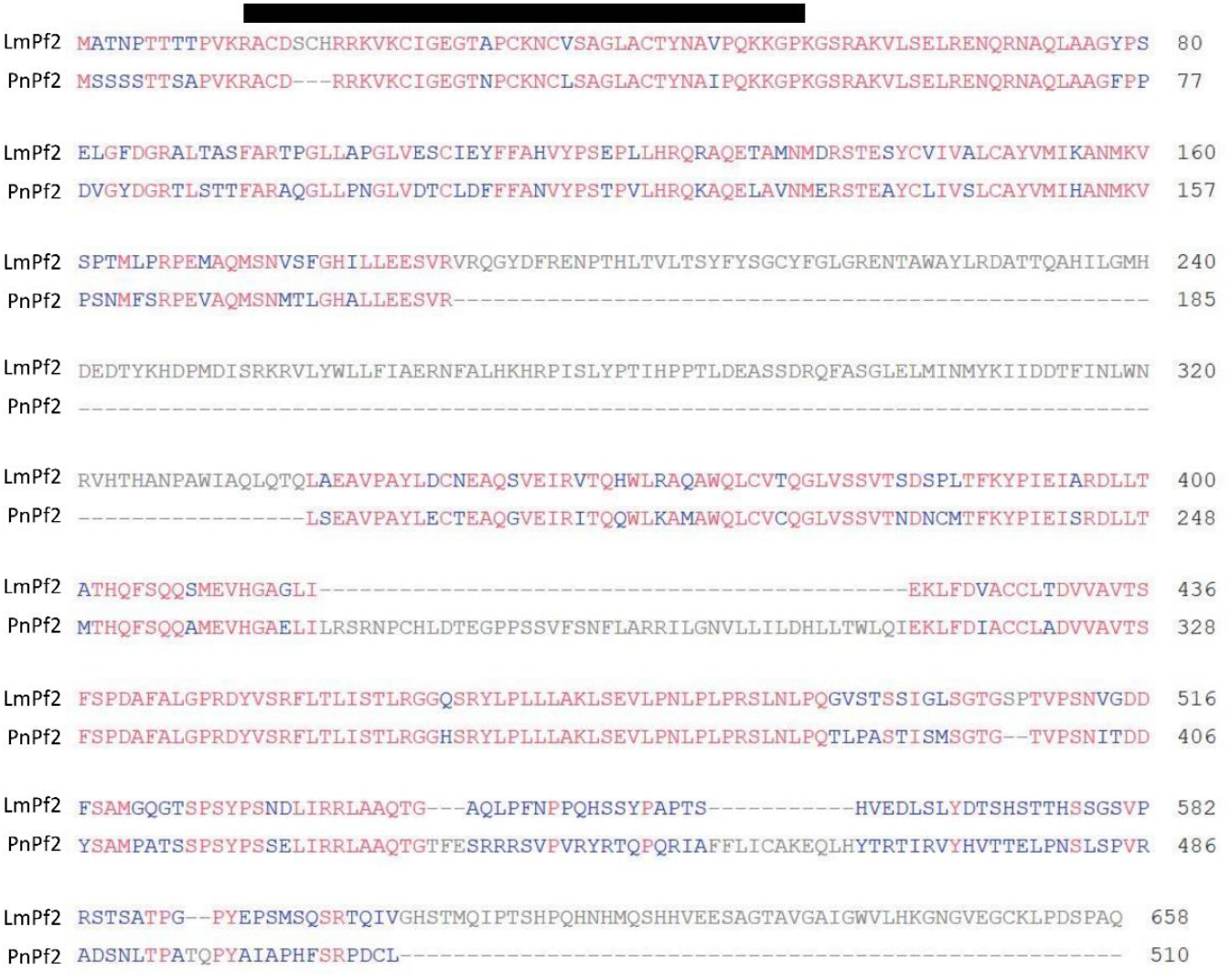
Identification of the PnPf2 orthologue in *Leptosphaeria maculans*. Alignment between LmPf2 from *Leptosphaeria maculans* and PnPf2 from *Parastagonospora nodorum* was performed using COBALT (Papadopoulos and Agarwala, 2007). The black bar corresponds to the Zn2Cys6-type DNA binding domain identified using Pfam (Finn *et al*., 2014; https://pfam.xfam.org/). LmPf2 and PnPf2 share 67% identity.

In a previous analysis, we identified the gene encoding KMT1 in *L. maculans* (gene ID Lmb_jn3_09141; Soyer *et al*., 2014; Dutreux *et al*., 2018). We investigated the expression profile of *LmPf2* and *KMT1* during axenic growth and at different stages of oilseed rape infection in controlled conditions and compared their expression profiles to that of the avirulence genes of *L. maculans* (**Figure 2**). As previously described, expression of *L. maculans* avirulence genes was repressed during axenic growth. During cotyledon infection, their expression increased strongly during the asymptomatic stage (between two and nine dpi), culminating at seven dpi and then, expression slowly decreased with concomitant appearance of necrotic symptoms (between 12 and 15 dpi; **Figure 2**). Likewise, *LmPf2* was not expressed during axenic growth while its expression was high at 2 dpi, peaked at 7 dpi and decreased until 15 dpi. *KMT1* was inversely expressed with a high expression during axenic growth, no expression during early infection (between two and nine dpi), and was up-regulated at 12 dpi when avirulence gene expression decreased. In contrast, during petiole and stem infection, avirulence genes and *LmPf2* were highly expressed during the asymptomatic growth in petioles (7 dpi) and in stems (14 and 28 dpi), while at these stages, *KMT1* was not expressed. To summarize, *LmPf2* showed an expression profile similar to that of the *L. maculans* avirulence genes, with an earlier induction of its expression compared to avirulence genes, while expression of *KMT1* was high during axenic growth and at late stages of cotyledon infection, when expression of avirulence genes is low (**Figure 2**).

**Figure 2:**
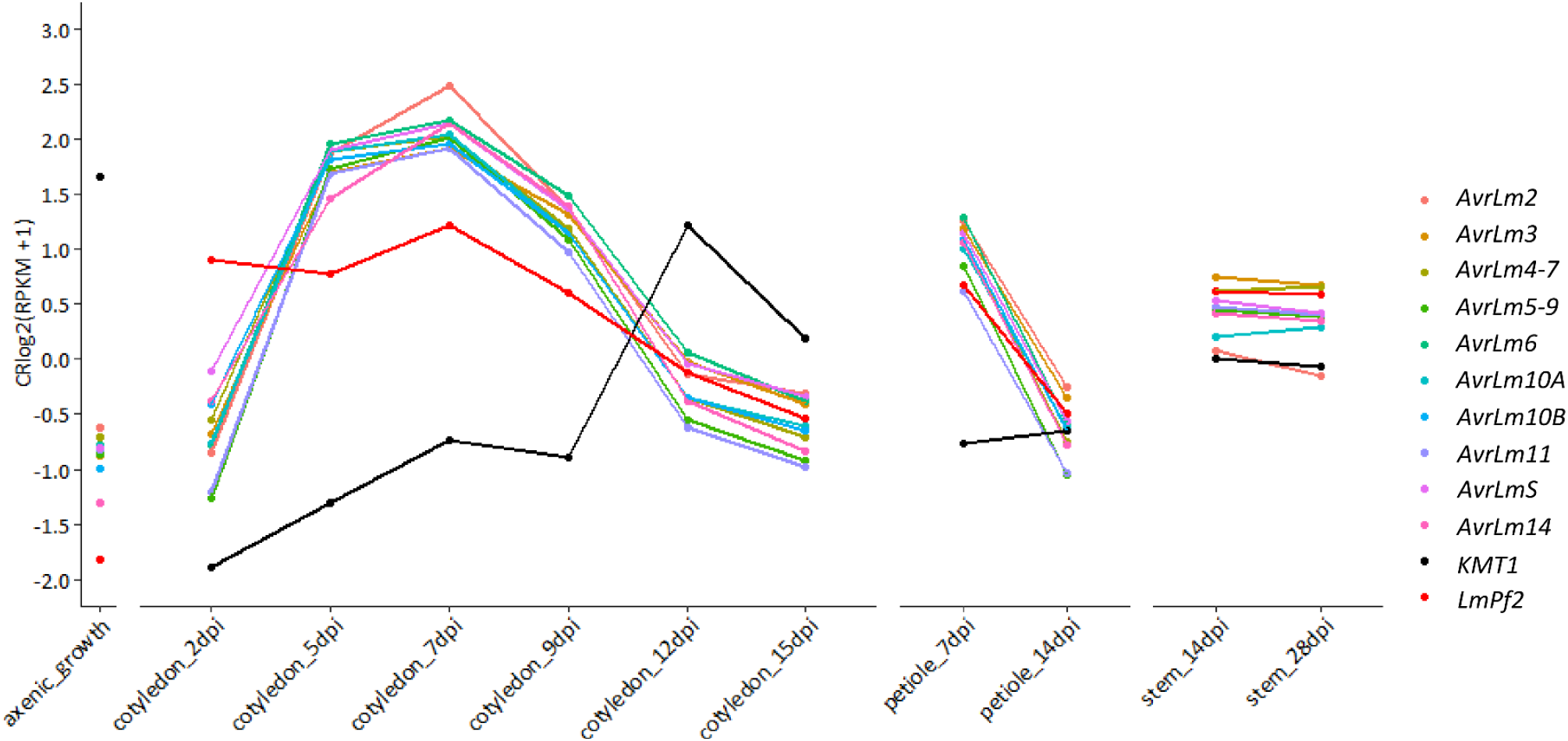
Expression profile of *Leptosphaeria maculans* avirulence genes, the *LmPf2* transcription factor and *KMT1* during axenic growth, infection of oilseed rape cotyledons, petioles and stem. Mycelium was obtained by growing the WT strain on V8-agar medium for 7 days (axenic growth). Oilseed rape was inoculated by the WT strain and sampled at different time points and on different organs (cotyledons at 2, 5, 7, 9, 12 and 15 days post infection, petiole at 7 and 14 dpi and stem at 14 and 28 dpi). The expression level of 10 avirulence genes, *LmPf2* and *KMT1* is represented by the log2 of RPKM (Reads Per Kilobase Per Million mapped reads) centered and reduced (Gay *et al*., 2021).

### Inactivation of *LmPf2* and *KMT1* do not induce morphological, conidiation or pigmentation defects while *LmPf2* over-expression induces developmental defects

Inactivation of *LmPf2* was performed using the CRISPR-Cas9 strategy and 16 transformants resistant to both hygromycin and geneticin were obtained and sequenced for the *LmPf2* gene. Among the 16 transformants, 12 had no mutations in *LmPf2* compared to the WT, three displayed a 1-bp deletion and one had a 5-bp deletion near the cleavage site (**Figure 3A**). These mutations resulted, at the protein level, in frame-shifts leading to two different truncated proteins of 175 amino-acids and 238 amino-acids respectively for the 1-bp and the 5-bp deletions compared to the 658 amino-acids length of the WT protein (**Figures 3B, C**). The two mutants were crossed with the WT strain constitutively expressing GFP (Materials and Methods) in order to select in the progeny *ΔLmPf2* and *ΔLmPf2*-*GFP* mutants without *Cas9* and CRISPR gRNA (**Table S2**). The four corresponding progeny strains are hereinafter referred to as *ΔLmPf2_*A, *ΔLmPf2_*A*-GFP, ΔLmPf2_*B *and ΔLmPf2_*B*-GFP* (_A corresponding to the 1-bp and _B to the 5-bp deletion). The four *ΔLmPf2* mutants were able to produce conidia (**Table 1** and data not shown) and their hyphae showed the same dark coloration similar to the WT after 14 days of growth on V8-juice agar medium (**Figure S1** and data not shown). After nine days of growth on V8-agar plate, only *ΔLmPf2_*B showed a significantly higher growth rate than the WT (Kruskal Wallis, *P value* <0.05; **Table1**) indicating that *LmPf2* inactivation did not lead to growth defect. To summarize, no major defect in conidia production, growth rate or morphology was associated with the inactivation of *LmPf2*.

**Table 1:** Influence of KMT1 and LmPf2 on axenic growth, conidia production, pathogenicity and effector gene expression in *Leptosphaeria maculans*.

| Isolate / transformants<sup>a</sup> | Growth 9 days post inoculation on V8 medium (mm)<sup>b</sup> | conidia/ml | Effector gene expression during axenic growth<sup>c</sup> |  |
| --- | --- | --- | --- | --- |
|  |  |  | *AvrLm6* | *AvrLm4-7* |
| WT | 60.5 (± 2.24) | 4.50.10<sup>7</sup> | 1 | 1 |
| *Δkmt1* | 50.25 (± 1.94) | 4.00.10<sup>7</sup> | 9.70.10<sup>-2</sup> (± 0.22) | 2.37.10<sup>2</sup> (± 27) |
| *ΔLmPf2_A* | 56.13 (± 0.9) | 1.53.10<sup>8</sup> | 4.60.10<sup>-2</sup> (± 0) | 0 |
| *ΔLmPf2_B* | 80 (± 0.8)* | 1.08.10<sup>8</sup> | 6.30.10<sup>-3</sup> (± 0) | 4.10.10<sup>-3</sup> (± 0) |
| WT_oPf2_A | 77.38 (± 0.76) | 0 | 3.05 (± 0.14) | 2.50.10<sup>-1</sup> (± 0.20) |
| WT_oPf2_B | 73.5 (± 1.19) | 0 | 2.68 (± 0.19) | 7.00.10<sup>-2</sup> (± 0.02) |
| *Δkmt1_oPf2_A* | 68.38 (± 1.03) | 2.00.10<sup>7</sup> | 2.61.10<sup>1</sup> (± 3.4) | 4.73.10<sup>2</sup> (± 8.6) |
| *Δkmt1_oPf2_B* | 53.75 (± 1.06) | 1.25.10<sup>6</sup> | 1.30.10<sup>4</sup> (± 107.79) | 2.31.10<sup>3</sup> (± 24.64) |

<sup>a</sup>*ΔLmPf2\_A* and *\_B* are two *LmPf2* mutants; WT\_oPf2\_A and WT\_oPf2\_B correspond to *LmPf2* over-expressing transformants in a WT background (with respectively a 600-fold and 150-fold over-expression of *LmPf2* compared to the WT); *Δkmt1\_oPf2\_A* and *Δkmt1\_oPf2\_B* correspond to *LmPf2* over-expressing transformants in a *Δkmt1* mutant background (with respectively a 150-fold and 600-fold expression of *LmPf2* compared to the *Δkmt1* mutant);
<sup>b</sup>Growth was monitored by measuring the diameter of the colony on the Petri dish;
<sup>c</sup>Mycelium was obtained by growing the WT strain and transformants in Fries liquid medium for 7 days. Two biological replicates per condition were generated. Total RNA was extracted. Expression of *AvrLm6* and *AvrLm4-7* was measured by qRT-PCR and expressed relatively to *LmβTubulin* expression and to expression of *AvrLm6* or *AvrLm4-7* in the WT using the 2<sup>-ΔΔCt</sup> method (Livak and Schmittgen, 2001);
\*: transformant statistically different from the WT, Kruskal Wallis test, *P-value* < 0.05.

**Figure 3:**
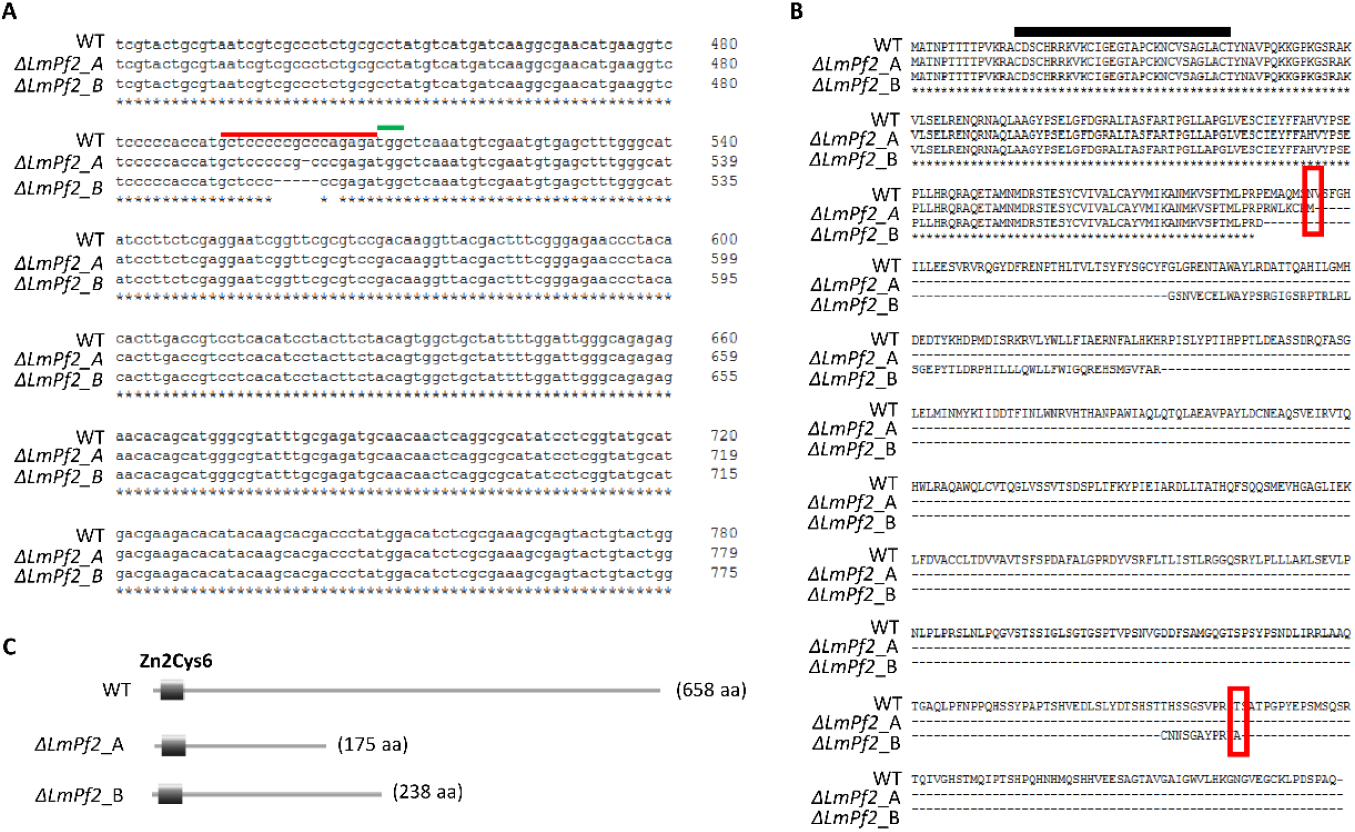
Effect of the *LmPf2* mutations on LmPf2 protein sequence. **A.** Alignment of the *LmPf2* gene of the WT isolate and two *LmPf2* mutants, *ΔLmPf2_A* and _B, showing respectively a 1-bp and 5-bp deletion. The PAM (Protospacer Adjacent Motif) is highlighted in green and the region targeted by the guide RNA is highlighted in red. **B.** Protein sequence of LmPf2 in the WT isolate and in the two *ΔLmPf2* mutants. The DNA-Binding Domain is indicated by a black bar. The red frames indicate the location of the stop codons in the mutant versions of the protein. **C.** LmPf2 protein length and domains identified with Pfam as described (Finn *et al*., 2014).

In a previous study, Soyer *et al*. (2014) silenced expression of *KMT1* (with a residual expression of 16% compared to the WT strain). Silencing of *KMT1* led to an over-expression of effector genes located in TE-rich regions, notably avirulence genes, during axenic growth. This over-expression was associated, at least for two avirulence genes, with a decrease of H3K9me3 at their loci. Here, we took advantage of the availability of the CRISPR-Cas9 strategy to better investigate involvement of KMT1 in the regulation of *L. maculans* gene expression. Twenty-five transformants resistant to hygromycin and geneticin were obtained and sequenced for the *KMT1* gene. Among the 25 transformants, 24 had no mutations in *KMT1* compared to the WT and one had a 1-bp insertion resulting, at the protein level, in a truncated protein of 144 aa (while the WT protein had a length of 516 amino-acids; **Figure 4A and B**). Both functional domains of KMT1 (Pre-SET and SET) were absent from the truncated protein (Pfam analysis; Finn *et al*., 2014; https://pfam.xfam.org/; **Figure 4C**) resulting in loss of H3K9me3 in the *Δkmt1* mutant strain confirmed by Western blot analysis (**Figure 4D**; **Figure S2**). The *Δkmt1* mutant was crossed with a WT-GFP strain in order to select in the progeny *Δkmt1* and *Δkmt1*-GFP mutants without *Cas9* and CRISPR gRNA (**Table S2**). The two selected progeny isolates are hereinafter referred to as *Δkmt1* and *Δkmt1-GFP* mutants. *Δkmt1* was not significantly altered in its axenic growth, morphology or conidia production (**Table 1**; **Figure S1** and data not shown).

**Figure 4:**
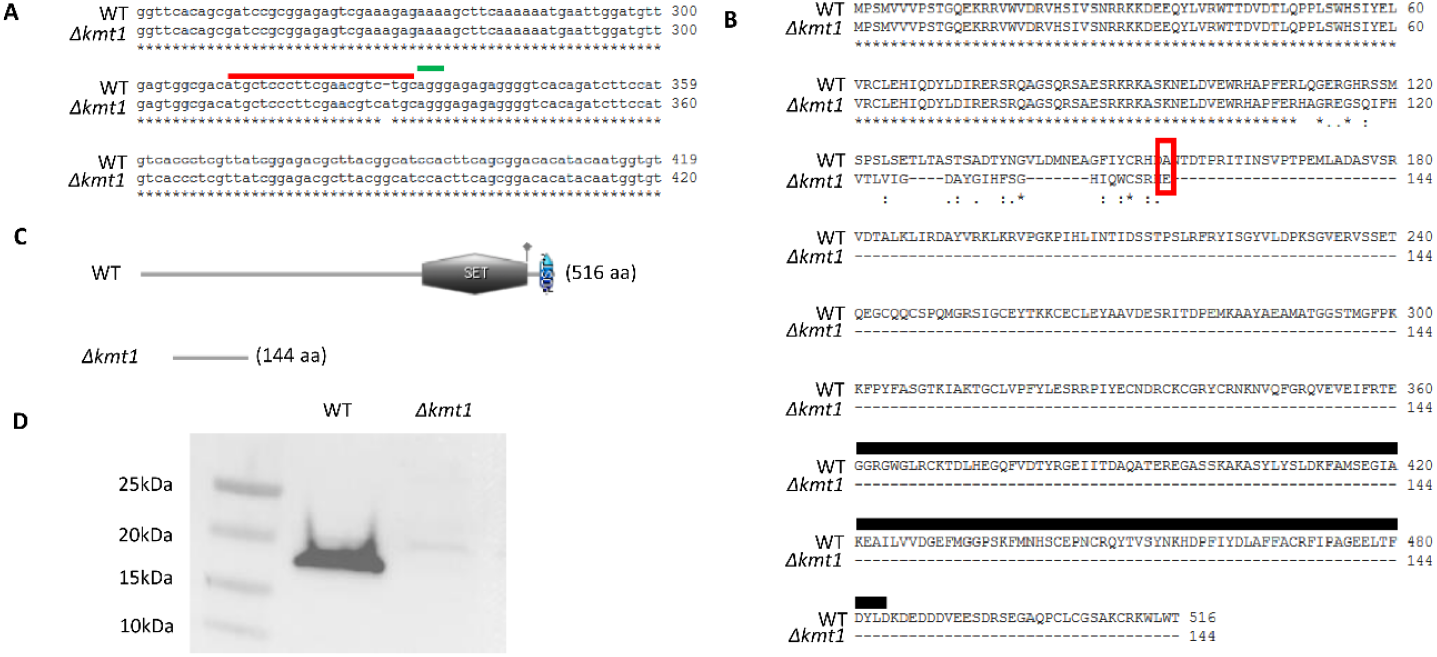
Effect of the *kmt1* mutation on the KMT1 protein sequence and function. **A.** Sequence alignment of the *KMT1* gene in the WT isolate and in the *Δkmt1* mutant showing a 1-bp insertion. The PAM (Protospacer Adjacent Motif) is highlighted in green and the region targeted by the guide RNA is highlighted in red. **B.** Protein sequence of KMT1 in the WT isolate and in the *Δkmt1* mutant. The red frame indicates the stop in the mutant version of the protein. The black bar indicates the SET domain which has been lost in the truncated protein. **C.** KMT1 protein length and domains identified with Pfam (Finn *et al*., 2014). **D.** Western Blot analysis of H3K9 tri-methylation in the WT isolate and in the *Δkmt1* mutant using an anti-H3K9me3 antibody (39062 Active Motif; see also Figure S2).

As association of effector genes enriched in H3K9me3-domains (deposited by KMT1) during axenic culture inhibits their expression, and that removal of H3K9me3 *in planta* might be a pre-requisite for concerted expression of effector genes (Soyer *et al*., 2014; 2015a; 2021), we analyzed effect of *LmPf2* over-expression in two different genetic backgrounds: in the WT isolate and in the *Δkmt1* mutant. Twenty-five transformants were recovered for each transformation and named respectively WT_o*Pf2* and *Δkmt1_*o*Pf2*. We measured expression of *LmPf2* during axenic growth by qRT-PCR in the transformants. Among the transformants, eight WT_o*Pf2* transformants showed a 147 to 18,000-fold increase of expression of *LmPf2* compared to the WT while *LmPf2* expression increased 108 to 596-fold in seven *Δkmt1_*o*Pf2* transformants (**Figure S3**). For further analyses, we selected two transformants from each genetic background with similar *LmPf2* expression levels (150 and 600 times more expressed than in the WT isolate or the *Δkmt1* transformant; **Figure S3**): WT_o*Pf2*_14, WT_o*Pf2*_24, *Δkmt1*_o*Pf2*_8 and *Δkmt1*_o*Pf2*_22 (hereafter referred to as WT_o*Pf2*_A, WT_o*Pf*2_B, *Δkmt1_*o*Pf2_*A and *Δkmt1_*o*Pf2_*B). Over-expression of *LmPf2* did not induce any growth defect even though the thallus was denser and harbored a white coloration. Over-expression of *LmPf2* had a critical impact on conidiation excepted for the *Δkmt1_*o*Pf2_*A transformant which was able to produce conidia but to a lesser extent than the WT isolate (**Table 1**; **Figure S1**).

### LmPf2 and KMT1 are involved in the pathogenicity of *L. maculans*

We inoculated the *ΔLmPf2, ΔLmPf2-GFP, Δkmt1, Δkmt1-GFP, LmPf2*-overexpressing transformants and the WT strain on cotyledons of a susceptible cultivar of oilseed rape. *Δkmt1* and *Δkmt1-GFP* showed reduced symptoms compared to the WT, indicating a decrease of pathogenicity (**Figure 5A-C; Figure S4**). Cotyledons infected with *Δkmt1-GFP* were observed from 5 to 13 dpi, which allowed us to distinguish living plant cells, fungal hyphae, and production of pycnidia. The *Δkmt1-GFP* transformant was able to colonize the plant and to produce pycnidia but induced less symptoms than the WT (**Figure 5; Figure S4**). The *ΔLmPf2* mutants were not able to invade the cotyledon further than the inoculation site, and consequently, did not induce any visible symptom (**Figure 5; Figure S4; Table 1**). Altogether, our results indicate that both KMT1 and LmPf2 are involved in infection establishment.

**Figure 5:**
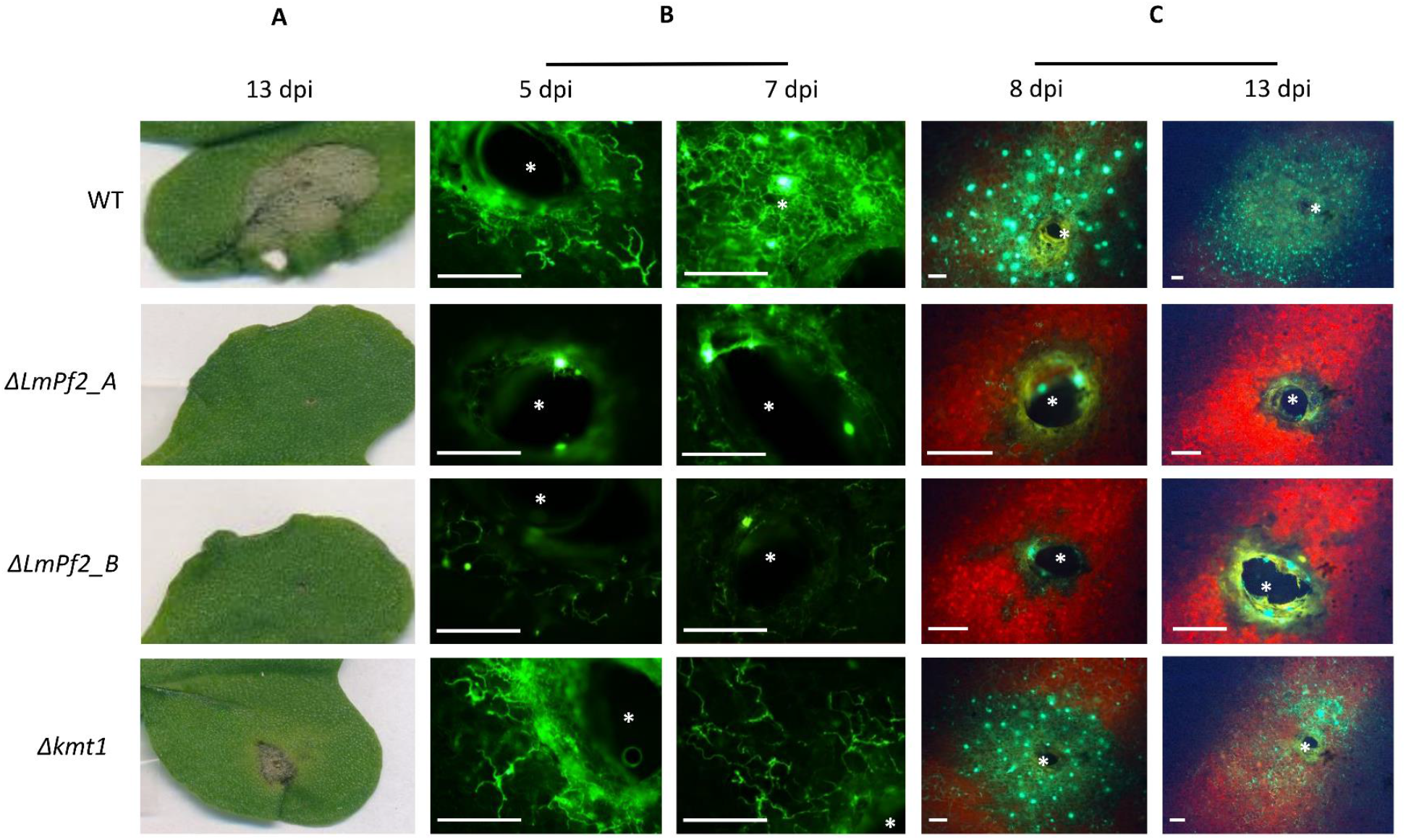
Effect of *LmPf2* and *kmt1* mutations on pathogenicity of *Leptosphaeria maculans*. The WT isolate and mutants inactivated for *LmPf2* or *KMT1* were inoculated on cotyledons of the susceptible cultivar of oilseed rape Es-Astrid. **A.** Pictures of symptoms at 13 dpi. **B.** Fluorescence images of cotyledons infected by GFP mutants and the WT isolate at 5 and 7 dpi using a confocal microscope. **C.** Fluorescence images of cotyledons infected by GFP mutants and the WT isolate at 8 and 13 dpi using a fluorescence binocular. Infection site (needle hole) is indicated by a white asterisk. Scale bars represent 1 mm. dpi = days post inoculation.

### LmPf2 and KMT1 are involved in the control of genes encoding known avirulence effectors

We investigated expression of two avirulence genes (*AvrLm4-7* and *AvrLm6*) in the *Δkmt1*, the *ΔLmPf2* mutants and in the WT strain during axenic growth by qRT-PCR (**Table 1**). During axenic growth, these genes are lowly expressed (Rouxel *et al*., 2011). In the *Δkmt1* mutant, *AvrLm6* was lowly expressed during axenic growth. On the contrary, *AvrLm4-7* was strongly over-expressed compared to the WT strain (237-fold more expressed; **Table 1**). *AvrLm4-7* and *AvrLm6* were lowly expressed in the *ΔLmPf2* mutants during the axenic culture, which is similar to expression of these genes in the WT strain (**Table 1**).

We also compared expression of *AvrLm4-7* and *AvrLm6* in the transformants over-expressing *LmPf2* (either in a WT or a *Δkmt1* background) and in the WT strain. Expression of *AvrLm6* slightly increased in the WT_o*Pf2* mutants (with a maximum of 3-fold increase compared to the WT) while *AvrLm4-7* was less expressed in the WT_o*Pf2* transformants than in the WT (Table 1). Over-expression of *LmPf2* in a *Δkmt1* background induced a huge increase of expression of both *AvrLm4-7* and *AvrLm6* during axenic culture (**Table 1**; with a maximum of 2,300- and 13,000-fold increase respectively for *AvrLm4-7* and *AvrLm6*). Noticeably, the level of over-expression of the *AvrLm* genes correlated with the level of overexpression of *Lmpf2*. We then hypothesized that an even stronger over-expression of *LmPf2* would bypass the negative regulatory effect of KMT1, to investigate whether removal of H3K9me3 would not be a pre-requisite for effector gene induction *in planta*. Hence, we investigated expression of *AvrLm4-7* and *AvrLm6* in two WT_o*Pf2* transformants (WT_o*Pf2*_22 and WT_o*Pf2*_25) in which *LmPf2* was over-expressed 10,000 and 18,000 times compared to the WT strain (**Figure S3, S5**). The strong *LmPf2* over-expression induced a higher up-regulation of *AvrLm4-7* and *AvrLm6* expression during axenic culture (with a maximum of 1,000-fold increase for *AvrLm4-7* and of 5,000-fold increase for *AvrLm6*). Nevertheless, it did not reach the same level as obtained when over-expressing *LmPf2* in a *Δkmt1* background (**Table 1 and Figure S5**).

We then investigated expression of four avirulence genes (*AvrLm4-7, AvrLm6, AvrLm10A* and *AvrLm11*) in the *ΔLmPf2*, the *Δkmt1* mutants and the WT strain during infection of oilseed rape at 7 dpi (when expression of *L. maculans* effectors reaches a peak in the WT strain). Inactivation of *LmPf2* strongly decreases expression of the four avirulence genes (1,000 to 2,000-fold less expressed in the *ΔLmPf2* mutant than in the WT) or even abolishes their expression (**Table 2**). Inactivation of *KMT1* also decreased, but to a lower extend, expression of the avirulence gene tested by qRT-PCR *in planta* (**Table 2)**. Altogether, our results confirm that KMT1 represses avirulence gene expression during axenic culture, suggest that LmPf2 positively regulates their expression and that KMT1 and LmPf2 act antagonistically to regulate expression of avirulence genes.

**Table 2:** Influence of LmPf2 and KMT1 on avirulence gene expression during infection of oilseed rape cotyledons.

| Isolate/<br>transformant | expression *in planta* 7 dpi<sup>a</sup> |  |  |  |
| --- | --- | --- | --- | --- |
|  | *AvrLm4-7* | *AvrLm6* | *AvrLm10A* | *AvrLm11* |
| WT | 3.25.10<sup>1</sup> | 7.41.10<sup>1</sup> | 2.92.10<sup>1</sup> | 1.96.10<sup>2</sup> |
| ΔLmPf2_A | nd | 4.58.10<sup>-2</sup> | 2.49.10<sup>-2</sup> | 7.64.10<sup>-2</sup> |
| ΔLmPf2_B | 4.08.10<sup>-3</sup> | 6.27.10<sup>-3</sup> | 3.25.10<sup>-2</sup> | 9.55.10<sup>-2</sup> |
| Δkmt1 | 1.03 | 4.23 | 1.93 | 5.23 |

Expression of *AvrLm4-7*, *AvrLm6*, *AvrLm10A* and *AvrLm11* at 7 dpi of cotyledons of the susceptible cultivar Es-Astrid in the ΔLmPf2 and Δkmt1 mutants and the WT isolate JN3.
<sup>a</sup>Gene expression levels are relative to *$\beta$ -tubulin* and calculated as described by Muller *et al.* (2002). Each value is the average of two biological replicates (two extractions from different biological replicates) and two technical replicates (two RT-PCR). dpi: days post inoculation.
nd: not detected.

### LmPf2 regulates sugar metabolism and CAZYme expression independently of the chromatin context

To further investigate involvement of LmPf2 and KMT1 in the regulation of *L. maculans* genes, notably effector genes, we performed RNA-seq analyses during axenic growth of inactivated mutants, over-expressing transformants (with similar *LmPf2* transcript level) and the WT strain (i.e., WT, *Δkmt1, ΔLmPf2_*A, *ΔLmPf2_*B, *WT_oPf2_*A, *WT_oPf2_*B, *Δkmt1_oPf2_*A and *Δkmt1_oPf2_*B; **Table S3**). Eighteen million reads were obtained, on average, for each sample. After pre-processing, between 72% and 88% paired-end reads were uniquely mapped, except for one technical replicate of the *ΔLmPf2_*A mutant for which only 39% of paired-end reads were mapped (**Table S3**). We then plotted the log2(RPKM+1) of each sample and observed that technical and biological replicates were consistent, except for the biological replicates of the *Δkmt1_oPf2* transformants (**Figure S6**). Based on that observation, data from biological replicates were merged for further analyses of gene expression, except for *Δkmt1_*o*Pf2_*A and *Δkmt1_*o*Pf2_*B that were considered separately for subsequent statistical analyses.

Among the different transformants, the transformant *Δkmt1* showed the fewest DEG compared to the WT strain, while the transformant *Δkmt1_*o*Pf2_*B had the highest number of DEG (272 and 1,792 genes respectively for *Δkmt1* and *Δkmt1_*o*Pf2_*B; **Table 3**). The higher number of DEG in the transformant *Δkmt1_*o*Pf2_*B than in the *Δkmt1_*o*Pf2_*A was consistent with the fact that level of expression of *LmPf2* was highest in the former (respectively 600 and 150-fold compared to the WT). In the genome of *L. maculans*, a GO annotation could be assigned to 5,076 genes of the 13,047 predicted genes (Dutreux *et al*., 2018). We set up an identification of the GO terms enriched between the different transformants and the WT strain to gain insight into the underlying metabolic processes influenced by any of the transformant generated. No GO enrichment was identified among the DEG in the *Δkmt1* or *ΔLmPf2* mutants compared to the WT. Genes up-regulated in the transformants over-expressing *LmPf2*, regardless of their genetic background, were enriched in GO categories involved in primary metabolism associated with carbohydrates uptake (e.g. starch, glucose, oligosaccharide metabolic process; **Tables S4, S5**). All seven GO categories enriched in the genes up-regulated due to over-expression of *LmPf2* in the *Δkmt1_*o*Pf2_*A transformant were also identified when over-expressing *LmPf2* in the *Δkmt1*_o*Pf2_*B transformant *Δkmt1*_o*Pf2_*B; **Table S5**). Thus, while *LmPf2* was considerably more over-expressed in *Δkmt1_*o*Pf2_*B than in *Δkmt1_*o*Pf2_*A and although they did not group together (**Figure S6**), over-expression of *LmPf2* influenced genes involved in similar processes. Seven GO categories were found enriched solely among genes up-regulated in *Δkmt1_*o*Pf2_*B (**Table S5**). No GO enrichment was identified for genes down-regulated in the *Δkmt1* mutant and the *Δkmt1_*o*Pf2_*A transformant. One GO category (GO:0055114) was detected as enriched in the down-regulated genes of the *ΔLmPf2* mutant, the WT_o*Pf2* and the *Δkmt1_*o*Pf2_*B transformants, encompassing genes involved in oxido-reduction processes (**Table S6**). Overall, our GO enrichment analysis suggests that CAZymes (i.e. enzymes involved in biosynthesis, metabolism and carbohydrate transport) were significantly regulated in the transformants over-expressing *LmPf2* in a *Δkmt1* background. In the genome of *L. maculans*, 330 genes are predicted as encoding CAZYmes (Dutreux *et al*., 2018) among which 109 genes were deregulated in at least one type of transformant generated in this study (**Table S7**). Altogether, our analysis confirmed that CAZymes were significantly regulated by LmPf2 and KMT1 (Chi^2^ test; *P value* < 0.05).

**Table 3:** Genes encoding effectors or located in particular chromatin domains differentially expressed after inactivation of *KMT1* or *LmPf2* or over-expression of *LmPf2*.

|  |  | Total genes | Effector genes | H3K4me2<br>genes<sup>a</sup> | H3K9me3<br>genes<sup>a</sup> | H3K27me3<br>genes<sup>a</sup> | H3K9me3/H3K27me3<br>genes<sup>a</sup> |
| --- | --- | --- | --- | --- | --- | --- | --- |
| Up-regulated | *Δkmt1* | 129 | 22* | 22<sup>#</sup> | 3 | 64* | 4* |
|  | *ΔLmpf2* | 202 | 20 | 68<sup>#</sup> | 4 | 57* | 5* |
|  | *WT_oPf2* | 247 | 46* | 46<sup>#</sup> | 2 | 101* | 6* |
|  | *Δkmt1_oPf2_A* | 275 | 59* | 38<sup>#</sup> | 11* | 144* | 15* |
|  | *Δkmt1_oPf2_B* | 931 | 154* | 233<sup>#</sup> | 25* | 329* | 24* |
| Down-regulated | *Δkmt1* | 143 | 27* | 35<sup>#</sup> | 10* | 42* | 9* |
|  | *ΔLmpf2* | 225 | 40* | 58<sup>#</sup> | 4 | 88* | 10* |
|  | *WT_oPf2* | 277 | 19 | 110<sup>#</sup> | 1 | 60* | 3 |
|  | *Δkmt1_oPf2_A* | 158 | 23* | 32<sup>#</sup> | 11* | 47* | 6* |
|  | *Δkmt1_oPf2_B* | 861 | 83 | 364<sup>#</sup> | 11 | 182* | 14* |

<sup>a</sup>Genes associated with H3K4me2 (di-methylation of lysine 4 of histone H3), H3K9me3 (tri-methylation of lysine 9 of histone H3), H3K27me3 (tri-methylation of lysine 27 of histone H3) or H3K9me3 and H3K27me3 during axenic growth of the WT isolate (Soyer *et al.*, 2021);
<sup>#</sup>genes statistically under-represented in a given category;
\*genes statistically enriched in a given category;
Statistical analyses were performed using Chi<sup>2</sup>, *P value* < 0.05.

### KMT1 and LmPf2 control expression of effector genes associated to H3K9me3 and H3K27me3 during axenic culture

In the genome of *L. maculans*, two types of heterochromatin domains have been identified, either associated with TE-rich genomic regions and H3K9me3 or associated with gene-rich regions in which H3K27me3-domains were detected. Both types of heterochromatin domains were enriched with effector genes (Soyer *et al*., 2021) and with genes significantly up-regulated in *planta* (Gay *et al*., 2021). We confronted the transcriptomic analyses of the different transformants generated in our study with previously generated ChIP-seq data to identify a possible effect of KMT1 or LmPf2 on the expression of genes associated either with eu- or heterochromatin modifications. We observed a strong effect of KMT1 and or LmPf2 on expression of genes located in H3K9me3-domains *in vitro* due to the global loss of this modification in the *Δkmt1* mutant (Figure 3; **Figure S2**), but also an effect of KMT1 and LmPf2 on expression of genes located in H3K27me3-domains (**Table 3**).We also investigated whether inactivation of *KMT1, LmPf2*, or over-expression of *LmPf2* influenced expression of pathogenicity-related genes predicted in *L. maculans* (SSP-encoding genes and genes involved in secondary metabolite biosynthesis). Among the 11 previously cloned avirulence genes of *L. maculans*, eight were deregulated in at least one of the transformants (**Table 4**; one avirulence gene, *AvrLm1*, was not present in the strain used as genetic background). Inactivation of *KMT1, LmPf2* or over-expression of *LmPf2* in the WT strain had little effect on expression of these genes (**Table 4**). On the contrary, over-expression of *LmPf2* in a *Δkmt1* background led to up-regulation of eight *AvrLm* genes compared to the WT during axenic culture (**Table 4**). Inactivation of *KMT1* or over-expression of *LmPf2* in a WT background had no effect on expression of *AvrLm* genes while these genes were induced due to over-expression of *LmPf2* in a *Δkmt1* background. In conclusion, RNA-seq data support our hypothesis that removal of H3K9me3 is a pre-requisite for induction of avirulence gene expression via action of the TF LmPf2 but also showed that LmPf2 may have specific target among effector genes.

**Table 4:** Influence of KMT1 and LmPf2 on expression of avirulence (*AvrLm*) genes, *KMT1* and *LmPf2* in *Leptosphaeria maculans* during axenic culture.

| ID | name | TE<sup>a</sup> | H3K4me2<sup>b</sup> | H3K9me3<sup>b</sup> | H3K27me3<sup>b</sup> | $\Delta kmt1^c$ | $\Delta LmPf2^c$ | WT_oPf2<sup>c</sup> | $\Delta kmt1\_oPf2\_A^c$ | $\Delta kmt1\_oPf2\_B^c$ |
| --- | --- | --- | --- | --- | --- | --- | --- | --- | --- | --- |
| Lmb_jn3_00001 | *AvrLm3* | yes | - | all_included | - | up | - | - | up | up |
| Lmb_jn3_03262 | *AvrLm4-7* | yes | - | all_included | - | - | - | - | up | up |
| Lmb_jn3_05547 | *AvrLm14* | yes | - | all_included | - | - | - | - | up | up |
| Lmb_jn3_06039 | *LmPf2* | - | overlap | - | all_included | - | down | up | up | up |
| Lmb_jn3_07862 | *AvrLm6* | yes | - | all_included | - | - | - | - | up | up |
| Lmb_jn3_07863 | *AvrLm2* | yes | - | 5'_included | - | down | - | - | - | up |
| Lmb_jn3_07874 | *AvrLm10_A* | yes | - | all_included | - | down | - | - | down | up |
| Lmb_jn3_07875 | *AvrLm10_B* | yes | - | all_included | all_included | - | down | - | up | up |
| Lmb_jn3_08343 | *AvrLmS-Lep2* | yes | - | all_included | - | - | - | - | up | up |
| Lmb_jn3_09141 | *KMT1* | - | 3'_included | - | overlap | - | - | - | - | - |
| Lmb_jn3_10106 | *AvrLm5-9* | yes | - | all_included | - | - | - | - | - | - |
| Lmb_jn3_12994 | *AvrLm11* | yes | - | all_included | - | - | - | - | - | - |
| Lmb_jn3_13126 | *AvrLm1\** | yes | - | all_included | - | N/A | N/A | N/A | N/A | N/A |

<sup>a</sup>Genes located in a TE-rich genomic context (Dutreux *et al.*, 2018);
<sup>b</sup>Genes associated with H3K4me2, H3K9me3 or H3K27me3 during axenic culture in the wild type strain (Soyer *et al.*, 2021);
<sup>c</sup>Genes deregulated in a given transformant compared to the WT strain during axenic culture (this analysis);
\**AvrLm1* is not present in the strain used for this RNA-seq analysis (i.e. strain JN2).

We then investigated whether we could observe the same effect on the 1,070 effector genes predicted in the genome of *L. maculans* (Gay *et al*., 2021). Genes up- or down-regulated in the *Δkmt1* mutant compared to the WT were enriched in effector genes (**Table 3**), confirming role of KMT1 in regulating expression of effector genes (Soyer *et al*., 2014). Considering all transformants in which *LmPf2* is over-expressed (either in a WT background or in a *Δkmt1* background), 185 effector genes (i.e. 17% of effector genes) were up-regulated (**Table S8**), with the transformant *Δkmt1_oPf2_B* exhibiting the largest number of up-regulated effector genes and the WT_o*Pf2* transformants having the lowest number of effector genes up-regulated (**Table 3; Table S8**). Finally, as *AvrLm* genes are all located within TE-rich environment of the *L. maculans* genome and associated with H3K9me3, except AvrLm10B associated with H3K9me3 and H3K27me3 (Rouxel *et al*., 2011; Soyer *et al*., 2021), we wanted to know whether removal of H3K9me3 in the *Δkmt1* mutant combined with over-expression of *LmPf2* would preferentially increase expression of effector genes associated with H3K9me3. As for avirulence genes, while inactivation of *KMT1* or *LmPf2*, or over-expression of *LmPf2* in a WT background resulted in up-regulation of respectively three or two effector genes located in a H3K9me3-domain, the over-expression of *LmPf2* in a *Δkmt1* background resulted in up-regulation of 20 out of the 36 effectors genes associated with H3K9me3 (**Table S9**). This analysis strengthens the hypothesis that KMT1 and the transcription factor LmPf2 work together, although through an opposite regulatory mechanism, to regulate the expression of effector genes localized in a H3K9me3 genomic context during axenic culture (including avirulence genes).

Finally, we investigated the influence of KMT1 and LmPf2 on expression of genes involved in secondary metabolism biosynthesis (PKS and NRPS). Among the 27 genes encoding NRPS or PKS (Dutreux *et al*., 2018), a maximum of three of them were up-regulated in the different transformants (three in the *Δkmt1_*o*Pf2_*B transformant; **Table S10**). In conclusion, NRPS or PKS-encoding genes did not appear to be regulated by KMT1 and / or LmPf2. Altogether, our analysis confirmed the key regulatory role of KMT1, highlighted the regulatory role of the transcription factor LmPf2 and suggested an antagonistic effect of both actors on the regulation of effector genes.

### LmPf2 and KMT1 influence expression of genes naturally over-expressed during infection of oilseed rape

We took advantage of the availability of transcriptomic data throughout the lifecycle of *L. maculans* on oilseed rape to investigate whether genes naturally up-regulated *in planta* were influenced by LmPf2 and / or KMT1. 1,207 genes were previously found up-regulated in at least one stage of the infection of oilseed rape compared to the axenic culture of *L. maculans* (Gay *et al*., 2021). Expression of genes up-regulated *in planta* were significantly regulated by LmPf2 or KMT1 as 31% were up-regulated in at least one of the transformants generated in this study (378 genes out of 1,207 genes; **Table S11**); (Chi^2^ test, *P* < 2.2.10^−16^). The largest number of up-regulated genes was observed in the *Δkmt1_*o*Pf2* transformants, making these genes significantly regulated by KMT1 and / or LmPf2. The effect was even stronger for effector-encoding genes, since, among 256 effector-encoding genes over-expressed *in planta*, 115 (45%) were up-regulated in at least one of the transformants, with again the largest number of up-regulated genes in the *Δkmt1_*o*Pf2_*B transformant (86 genes, 34%). Our analysis shows that KMT1 inhibits while LmPf2 positively regulates expression of genes naturally expressed *in planta*.

## DISCUSSION

Focusing at the time on cotyledon infection, Soyer *et al*. (2015) proposed a two-layer regulatory model in which expression of *L. maculans* effector genes located in repeat-rich regions was repressed *in vitro* through H3K9me3 deposition by KMT1. They hypothesized that, *in planta*, an unknown signal triggered chromatin remodeling in the genomic environment of these effector genes allowing the binding of one or several transcription factor(s) leading to a concerted expression of effector genes (and possibly other pathogenicity-related genes located in TE-rich environment) during the primary infection of oilseed rape. Our results include other stages of plant colonization (petioles and stems), confirm that KMT1 represses effector gene expression during axenic culture, and show that LmPf2 positively regulates their expression and that both KMT1 and LmPf2 act together, in an opposite manner, to concertedly regulate expression of effector genes. Notably, *LmPf2* has an expression profile similar to that of the *L. maculans* avirulence genes and effector genes expressed during the asymptomatic phases of infection, while expression of *KMT1* is inversely correlated during axenic growth, cotyledon and petiole infection. To our knowledge, this is the first evidence of a double control involving a repressive histone modification and a specific TF on expression of effector genes in a fungal species. Altogether, these results allowed us to refine the proposed model for the double control of effector gene expression mediated by KMT1 and LmPf2 in *L. maculans*.

### LmPf2 is involved in the establishment of infection by *L. maculans*

In this study, we investigated the function of LmPf2 in *L. maculans*. CRISPR-Cas9 inactivation of *LmPf2* induced pathogenicity defects, with a very limited development of the fungus at the inoculation site, while the mutants had no alteration in their axenic growth or conidiation. In contrast, overexpression of *LmPf2* led to sporulation defects even if the transformants still induced symptoms on oilseed rape when inoculated through mycelial plugs. We conclude that, in *L. maculans*, LmPf*2* is involved in the establishment of oilseed rape infection. In three other Pleosporales species, *i*.*e. A. brassicicola, P. tritici repentis* and *P. nodorum*, Pf2 was also essential for the establishment of infection (Cho *et al*., 2013; Rybak *et al*., 2017; Jones *et al*., 2019). However, in *A. brassicicola*, AbPf2 was dispensable for normal growth while crucial for virulence on various Brassicaceae species (Cho *et al*., 2013). In *P. tritici-repentis, Pf2* mutants were both altered in their virulence on susceptible wheat cultivars and in their axenic growth and conidiation (Rybak *et al*., 2017). In *Zymoseptoria tritici*, a Pf2 orthologue was essential for virulence, but also regulates dimorphic switch, axenic growth and fungal cell wall composition (Habig *et al*., 2020). So, while the involvement of Pf2 in pathogenicity is a common feature, its involvement in developmental processes or morphological switches is species-dependent (John *et al*., 2021).

### LmPf2 controls carbon acquisition and cell-wall integrity independently of the chromatin context

RNA-seq analyses in *A. brassicicola* and *P. nodorum* using *Pf2* knockout mutants suggested that Pf2 controls the expression of a wide range of CWDEs during early infection (Cho *et al*., 2013; Jones *et al*., 2019). PnPf2 positively regulates CAZymes, CWDEs, peptidases and hydrolases, while it negatively regulates general metabolic activity, possibly to conserve energy (Jones *et al*., 2019). In Z. *tritici*, the Pf2 orthologue regulates carbon-sensing pathways (Habig *et al*., 2020). In *L. maculans*, LmPf2 regulates sugar metabolism and CAZYme expression independently of the chromatin context. Conservation of Pf2 in many Pleosporales together with its involvement in regulation of CAZymes, CWDEs, peptidases and hydrolases suggest that a shared evolutionary origin exists in the regulation of carbon acquisition in Pleosporales. It also indicates that the function of Pf2 has been expanded in the course of evolution to the regulation of fungal effectors, in link with a chromatin-based control in *L. maculans* (see below).

### KMT1 is involved in fungal aggressiveness and in the control of effector gene expression in *L. maculans*

We also investigated function of KMT1 in *L. maculans. Δkmt1* mutants displayed reduced aggressiveness on oilseed rape but normal growth and conidiation. This result contrasts with data obtained in *N. crassa, A. fumigatus* and *Z. tritici* in which the inactivation of *KMT1* led to growth defects (Tamaru and Selker, 2001; Palmer *et al*., 2008; Möller *et al*., 2019), but confirms involvement of KMT1 in fungal pathogenicity found in *Z. tritici* (Möller *et al*., 2019). Histone modification enzymes, notably KMT1, are increasingly documented for their ability to control concerted expression of secondary metabolite gene clusters and effector genes showing distinct genomic locations (Gacek and Strauss, 2012; Soyer *et al*., 2015; Collemare and Seidl, 2019). In the fungal endophyte *Epichloë festucea*, KMT1 regulates synthesis of symbiosis-specific alkaloids, which act as bioprotective metabolites, and is crucial for establishment of mutualistic interaction (Chujo and Scott, 2014). In *L. maculans*, Soyer *et al*. (2014) found that partial silencing of *KMT1* through RNAi led to avirulence gene over-expression and H3K9me3 depletion (at least at two avirulence genes loci), and allowed up-regulation of 30% of the genes located in TE-rich regions during axenic culture. In this study, while inactivation of *KMT1* had a significant impact on effector gene expression, only one avirulence gene and three genes located in H3K9me3 domains were up-regulated *in vitro*. In contrast, inactivation of *KMT1* had a significant effect on genes located in H3K27me3 domains. We hypothesize that the complete inactivation of *KMT1* could have led to H3K27me3 relocation at native H3K9me3 domains, as previously reported for *N. crassa* or *Z. tritici* (Basenko *et al*., 2015; Möller *et al*., 2019). This phenomenon might be less pronounced when silencing *KMT1* as the gene was still expressed at almost 20%, suggesting that the KMT1 activity, hence H3K9me3 deposition, was not completely abolished in our transformants (Soyer *et al*., 2014). The data presented here, notably the wide effect on expression of genes encoding effectors due to over-expression of *LmPf2* in a *Δkmt1* background, nevertheless suggest that KMT1 is involved in the control of *L. maculans* effector gene expression and of genes located in heterochromatin regions.

### LmPf2 controls effector gene expression in a *Δkmt1* mutant background

In other Pleosporales, no investigation of a link between a histone-modifying enzyme and Pf2 was performed, and no over-expression of *Pf2* performed although it is demonstrated to be a positive regulator of effector genes. In this study, we highlighted a major role of LmPf2 in the control of effector gene expression in *L. maculans*. We determined that *LmPf2* inactivation led to effector gene expression defect *in planta*. Furthermore, *LmPf2* over-expression in a *Δkmt1* background significantly induces expression of i) up to 154 effector genes including eight avirulence genes, ii) 378 genes associated with heterochromatin, regardless of the nature of the encoded protein and iii) that up-regulation of avirulence genes was much higher when LmPf2 was over-expressed in a *Δkmt1* than in a WT background.. These results are consistent with previous studies that described Pf2 as a regulator of effector gene expression in three Pleosporales species (Cho *et al*., 2013; Rybak *et al*., 2017; Jones *et al*., 2019). Pf2 was reported to regulate expression of 33 genes encoding putative secreted proteins including eight putative effectors in *A. brassicicola* (Cho *et al*., 2013). In *P. nodorum*, PnPf2 was found to be an essential regulator of both *ToxA* and *Tox3* expression, but only moderately involved in *Tox1* regulation, while the orthologue of *SnToxA, ToxA*, was regulated by PtrPf2 in *P. tritici-repentis* (Rybak *et al*., 2017). RNA-seq analyses in *P. nodorum* using *PnPf2* knockout mutants also suggested that PnPf2 regulates a wide range of uncharacterized effector-like genes during early infection (Jones *et al*., 2019). Altogether, these studies pointed out Pf2 as an important positive regulator of effector gene expression. However, these studies only reported effect of *Pf2* inactivation on gene expression and no over-expression of *Pf2* was performed. In *L. maculans*, we have investigated involvement of LmPf2 on regulation of gene expression not only through its inactivation but also via its over-expression in two different background. While the information on the chromatin context of effector genes is generally unavailable for the above-mentioned examples, in *L. maculans*, we highlighted a major effect of the chromatin-context on the ability of LmPf2 to regulate effector gene expression. Whether that model of double control of effector gene expression involving a specific TF and a histone-modifying protein could be generalized to other pathogenic fungi or if it is specific of *L. maculans* need to be investigated. For instance, in *Z. tritici*, inactivation of *KMT6* led to effector gene up-regulation that did not reach the expression level during wheat infection, suggesting the additional involvement of transcription factor(s) (Meile *et al*., 2020). In contrast, in *F. oxysporum* f. sp. lycopersici, the transcription factors Sge1, FTF1 and FTF2 were able to regulate expression of effector genes located on a pathogenicity dispensable chromosome independently of chromatin-remodeling (van der Does *et al*., 2016).

## Conclusions

All of these data provide information to refine the model proposed by Soyer *et al*. (2015) and build an up-dated model. i) We firstly show that LmPf2 is a master regulator for expression of CAZymes, CWDEs, peptidases and hydrolases along with more than 500 genes associated with heterochromatin, and with a strong enrichment in effector genes, including most of the avirulence genes identified in *L. maculans*; ii) the regulation of *LmPf2* expression mirrors that of the genes included in the “biotrophy” wave, with rounds of up and down regulation; iii) the genes up-regulated during infection and associated with heterochromatin context, and notably, a H3K9me3 context, are less accessible to the TF as long as the chromatin is condensed. Accessibility during the asymptomatic stages of cotyledon infection is thus rendered possible by the reduced expression of *KMT1*; iv) at the end of the asymptomatic stage of colonization of cotyledons, set up of necrotrophy involves chromatin condensation due to *KMT1* expression, and reduced expression of *LmPf2*, resulting in extinction of expression of genes involved in the “biotrophy” wave; v) the process is repeated identically during asymptomatic colonization of petioles and stems. This indicates that, in *L. maculans*, most avirulence genes and a significant number of effector genes expressed during the asymptomatic stages of oilseed rape infection are under the double control of KMT1 and LmPf2

## ACKNOWLEDGEMENT

Authors wish to thank all members of the “Effectors and Pathogenesis of *L. maculans*” group, as well as Jonathan Grandaubert for fruitful discussion. We are grateful to the BIOGER bioinformatics platform (https://bioinfo.bioger.inrae.fr/; Nicolas Lapalu, Adeline Simon) for providing support and storage resources. We thank Alexander Idnurm (Melbourne University, Australia) for providing us with the vector and protocols for the CRISPR-Cas9 approach. C. Clairet and EJ. Gay were founded by a PhD salary from the University Paris-Saclay; A. Porquier was founded by the ANR-PRC Project CHOPIN (ANR-19-CE20-0022-01). This work was partly funded by the “Plant Health and Environment” division of INRAE (ChromaDyn Project). The “Effectors and Pathogenesis of *L. maculans*” group benefits from the support of Saclay Plant Sciences-SPS (ANR-17-EUR-0007).

