## Supplementary Figures for "Regulation of effector gene expression as concerted waves in *Leptosphaeria maculans*: a two-players game"

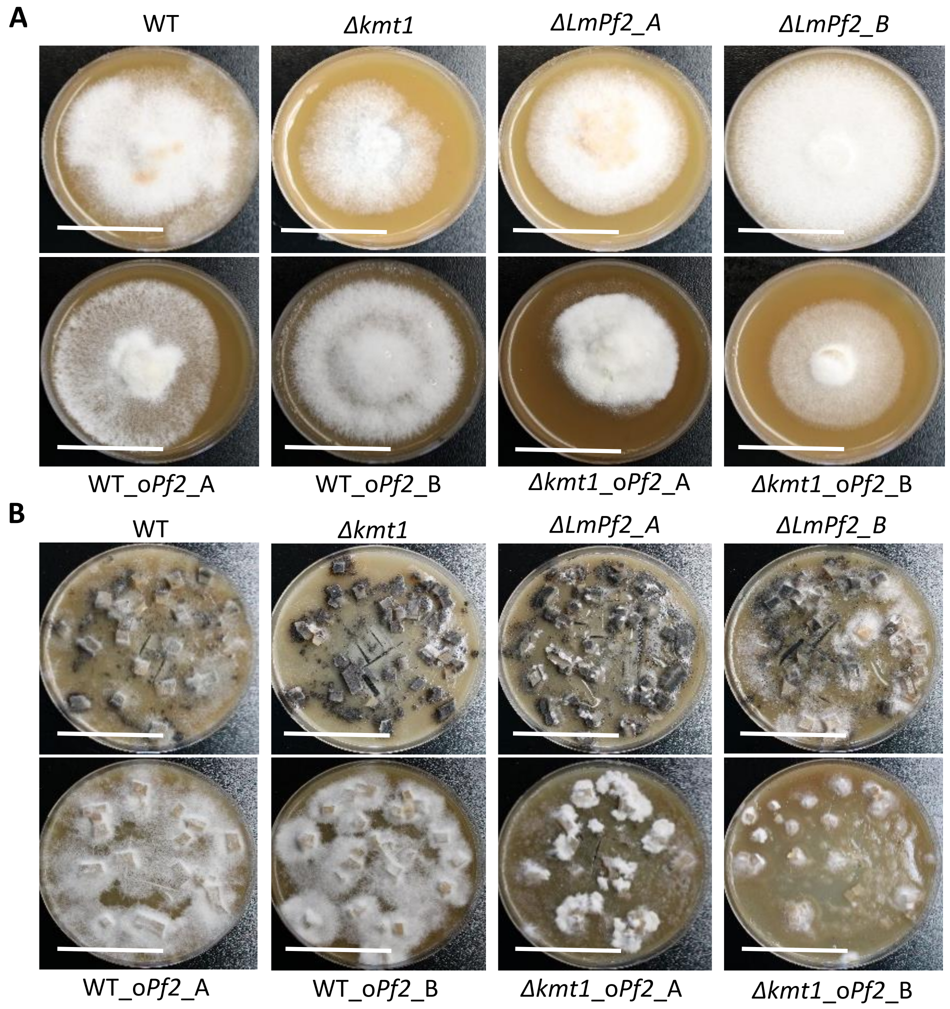


**Figure S1: Morphological characterization of *ΔLmPf2* mutants, *Δkmt1* mutants and transformants overexpressing *LmPf2*.** **A.** Transformant thallus morphology on V8 agar medium at 6 days after inoculation. Scale bars correspond to 25 mm. **B.** Transformant thallus morphology during conidiation on V8 agar 14 days after inoculation. Scale bars correspond to 45 mm. The transformants over-expressing *LmPf2* were obtained in a WT background and in a *Δkmt1* background.

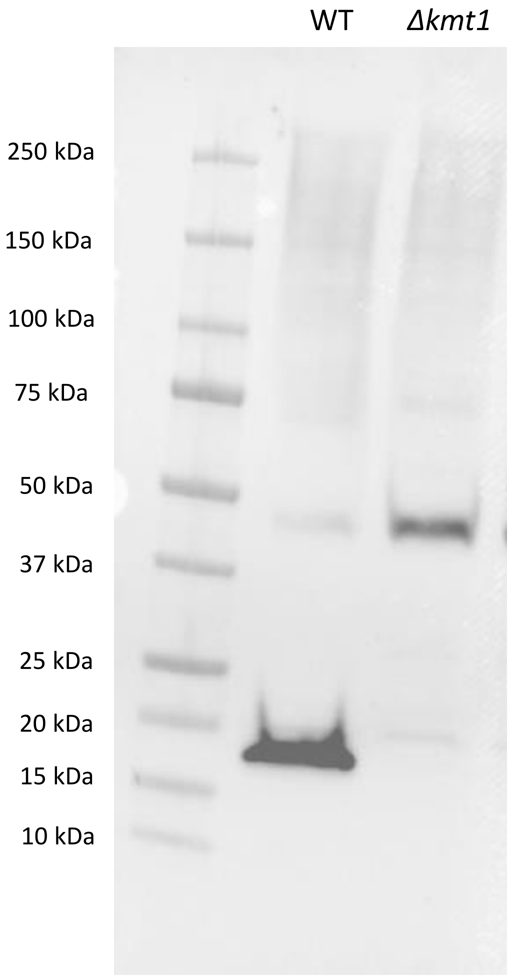


**Figure S2: Western Blot analysis of H3K9 tri-methylation in the WT isolate WT and *Δkmt1.*** Nuclear proteins were extracted from mycelium, separated by SDS-PAGE, blotted and probed with antibodies against H3K9me3.

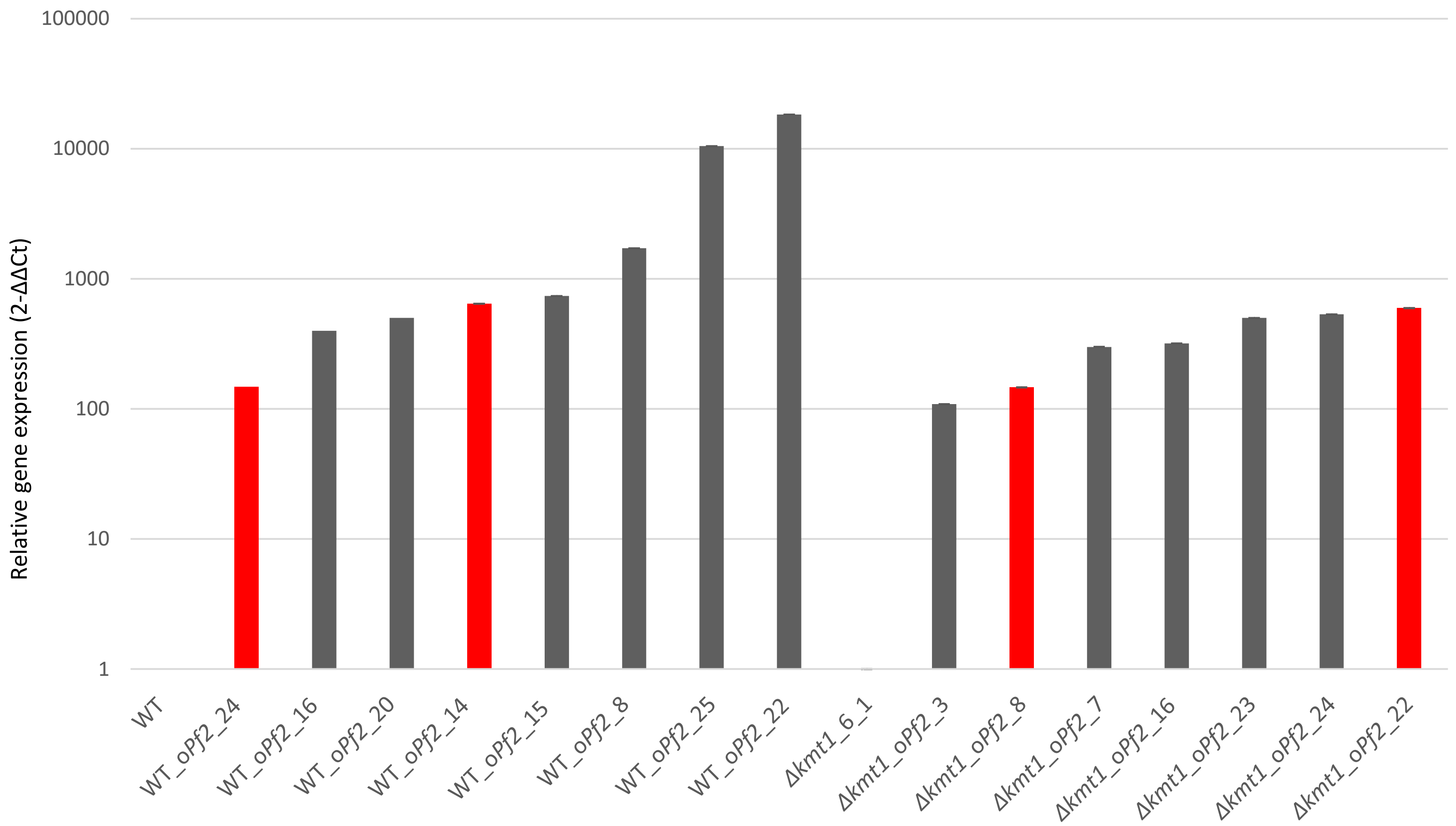


**Figure S3: Level of *LmPf2* expression in the *Leptosphaeria maculans* transformants over-expressing *LmPf2* during axenic growth.** Mycelium was obtained by growing WT strain and transformants in Fries liquid medium for 7 days. Two biological replicates per condition were generated. Total RNA was extracted. Expression of *LmPf2* was measured by qRT-PCR and expressed relatively to *LmβTubulin* expression and to expression of *LmPf2* in the strain before transformation using the $2^{-\Delta\Delta Ct}$ method (Livak and Schittgen, 2001). Red bars correspond to transformants which were selected for further characterization (WT_o*Pf2*_A, WT_o*Pf2*_B, *Δkmt1*_o*Pf2*_A and *Δkmt1*_o*Pf2*_B).

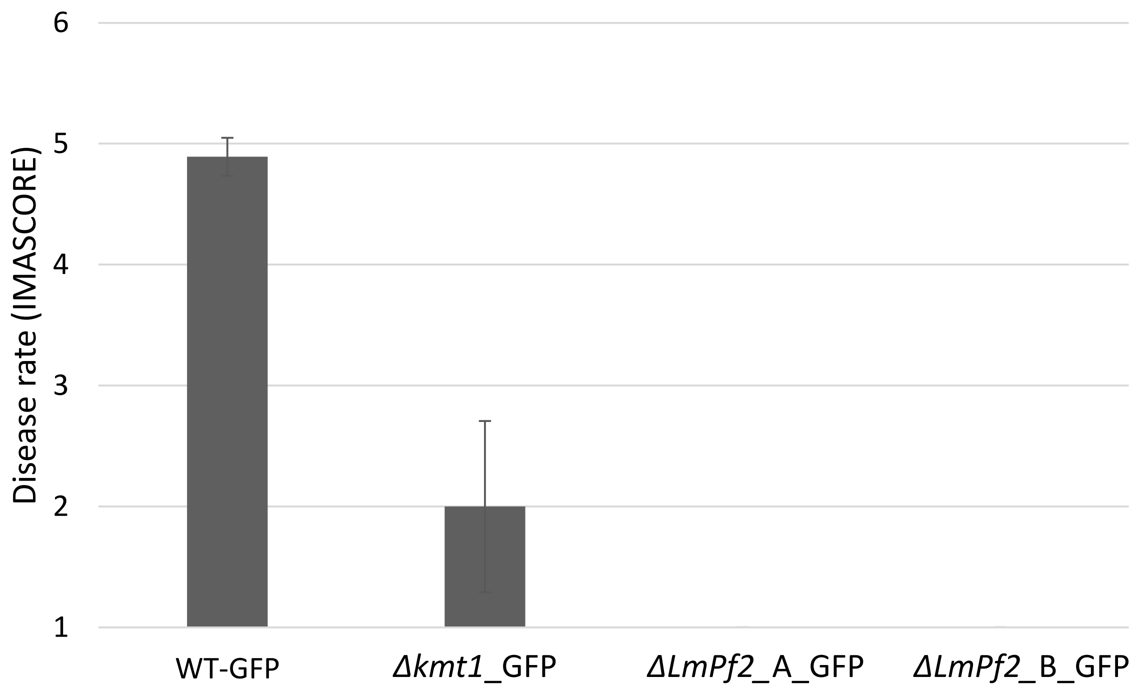


**Figure S4: Pathogenicity assays using a WT-GFP transformant, *Δkmt1* and *ΔLmPf2* transformants after crossing with WT-GFP.** Symptoms rating at 14 dpi on the susceptible cultivar of oilseed rape Es-Astrid after inoculation with three different CRISPR-Cas9 mutants and the WT-GFP transformant. Disease rate is expressed as the mean scoring using the IMASCORE rating scale comprising six infection classes (IC), where IC1 to IC3 correspond to various levels of resistance of the plant and IC4 to IC6 to susceptibility (Balesdent *et al.,* 2001).

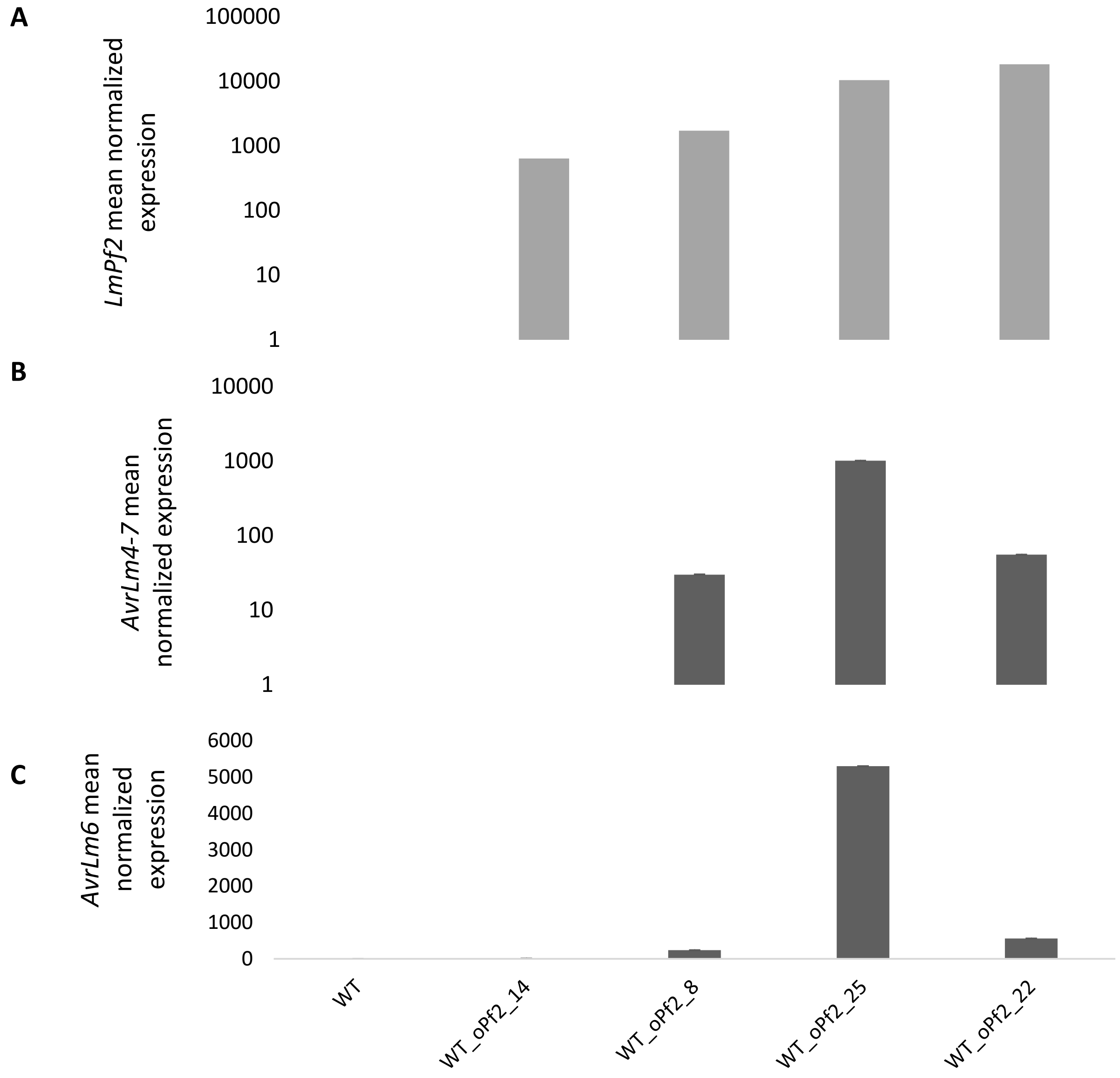


**Figure S5: Influence of *LmPf2* over-expression on the expression of two *L. maculans* avirulence genes during axenic growth.** Mycelium was grown in Fries liquid medium during seven days. Expression of **A.** *LmPf2*, **B.** *AvrLm4-7* and **C.** *AvrLm6* was measured by qRT-PCR using *β-Tubulin* as a constitutive reporter gene and expression in the WT as reference ($2^{-\Delta\Delta Ct}$ method). Each value is the average of two biological replicates (two extractions from different biological replicates) and two technical replicates (two RT-PCR).

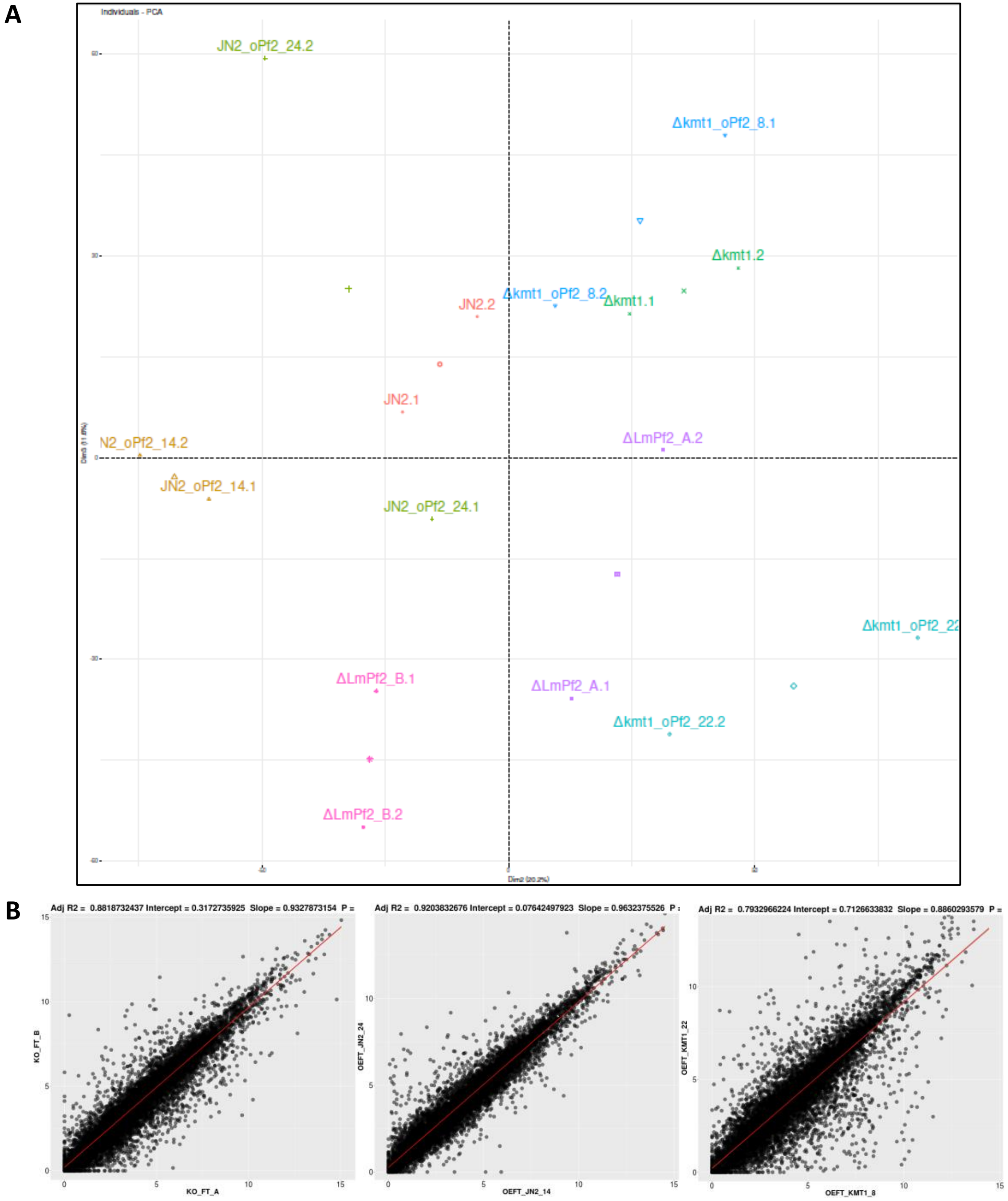


**Figure S6: Reproducibility of RNA-seq replicates. A.** log2(RPKM) PCA of all samples. Legend labels: WT: Wild type strain; Δ*kmt1*_X: Δ*kmt1_A* and _B; Δ*LmPf2*_X: Δ*LmPf2_A* and _B; WT_o*Pf2*_X: WT_o*Pf2*_A and _B; Δ*kmt1_oPf2*_X: *Δkmt1_oPf2*_A and _B. **B.** Scatterplot obtained with Log2(RPKM) of three biological replicates. Left panel: Δ*LmPf2_A* vs Δ*LmPf2_B;* middle panel: WT_o*Pf2*_A vs WT_o*Pf2*_B; right panel: *Δkmt1_oPf2*_A vs *Δkmt1_oPf2*_B.
